# Vasopressin-to-Oxytocin Receptor Crosstalk in the Preoptic Area Underlying Parental Behaviors in Male Mice

**DOI:** 10.1101/2024.07.01.601605

**Authors:** Kengo Inada, Mitsue Hagihara, Kasane Yaguchi, Satsuki Irie, Yukiko U. Inoue, Takayoshi Inoue, Kazunari Miyamichi

## Abstract

The transition to parenthood brings significant changes in behavior toward offspring. For instance, in anticipation of their offspring, male mice shift from infanticidal to caregiving behaviors. While the release of oxytocin from the paraventricular hypothalamus (PVH) plays a critical role in paternal caregiving, it does not fully account for the entire behavioral shift. The specific downstream neurons and signaling mechanisms involved in this process remain obscure. Here, we demonstrate that PVH vasopressin neurons also essentially contribute to a paternal behavioral shift. This vasopressin signal is partially transmitted through oxytocin receptors (OTRs) expressed in the anterior commissure and medial nuclei of the preoptic area. These OTR-expressing neurons receive inputs from both PVH oxytocin and vasopressin neurons and are responsible for expressing paternal caregiving behaviors. Collectively, this non-canonical vasopressin-to-OTR crosstalk within specific limbic circuits acts as a pivotal regulator of paternal behavioral changes in mice.

**Highlights:**

- PVH vasopressin neurons are required for and can trigger paternal caregiving behaviors.
- Vasopressin-induced paternal behaviors are mediated in part by OTRs in the preoptic area (POA).
- POA OTR neurons receive inputs from both PVH oxytocin and vasopressin neurons.
- POA OTR neurons play a critical and facilitative role in promoting paternal caregiving behaviors.

## Introduction

Across mammalian species, non-parental adult animals often show little interest in, or may even display aggressive behaviors toward, conspecific young. It is posited that infant-directed aggression (infanticide) has evolved in both males and females as a strategy to enhance their reproductive fitness by increasing mating opportunities and reallocating essential resources for future progeny^1,2^. The onset of caregiving behaviors toward infants is typically observed as animals anticipate their offspring^3,4^. Male laboratory mice have been a valuable model for investigating the behavioral shift associated with different life stages. While sexually naïve adult male mice are highly aggressive and may engage in infanticide, they undergo a shift toward caregiving behaviors upon mating and cohabitating with pregnant female mice^3,5,6^.

Studies in male rodents have identified specific limbic structures that play pivotal roles in infanticide or parental behaviors. For instance, the bed nucleus of the stria terminalis^7^, medial amygdala^8^, perifornical area of the hypothalamus^9^, and amygdalohippocampal area (AHi)^10^ harbor distinct neural populations responsible for male infanticide. By contrast, neurons expressing galanin^11,12^ or calcitonin receptor (Calcr)^13,14^ in the medial preoptic nucleus (MPN) act as positive regulators of paternal caregiving behaviors, in which prolactin receptor signaling is involved in activating galanin-positive neurons^15^. A recent study on maternal caregiving behavior proposed mutually suppressive antagonistic circuits between neurons promoting infanticide and those promoting caregiving behaviors^16^. However, whether similar mechanisms are utilized by male mice remains unclear. Overall, the molecular and neural mechanisms underlying paternal behavioral plasticity toward infants remain poorly understood.

Peptide hormones are thought to play a role in paternal behaviors^17^. For instance, biparental male mandarin voles display time-locked activity of paraventricular hypothalamus (PVH) neurons producing oxytocin (OT), a nonapeptide hormone, during paternal caregiving behaviors^18^. Additionally, our recent study in mice found that PVH OT neurons are crucial for regulating paternal caregiving behaviors^19^. Specifically, chemogenetic activation of PVH OT neurons effectively reduces infanticidal behaviors and promotes pup retrieval, a hallmark of parental behavior^20^, in sexually naïve male mice, with this effect being OT ligand-dependent. Conversely, conditional knockout (cKO) of the *OT* gene restricted to the PVH in adulthood results in a significant decrease in paternal caregiving behaviors, leading fathers to ignore pups without exhibiting attack or retrieval behaviors. These data suggest a model of paternal behavioral transition occurring in a stepwise manner, from infanticidal to ignoring (Step I) and then to parenting (Step II)^19^. While chemogenetic activation of PVH OT neurons can induce both Steps I and II, the loss-of-function of PVH OT signals still demonstrates Step I transition in fathers, suggesting the presence of compensatory mechanisms beyond OT. One potential candidate is arginine vasopressin (AVP), also known as mammalian vasotocin^21^, which is a closely related neural hormone involved in various social behaviors^22^. While a classical study found that intracerebroventricular (icv) administration of AVP facilitates maternal behaviors in virgin rats^23^, the specific role of PVH AVP neurons or the AVP ligands they release in paternal behaviors remains elusive.

OT has a single canonical receptor type known as the OT receptor (OTR), whereas AVP interacts with three AVP receptors (VRs) named V1aR, V1bR, and V2R^21,24^. In the mammalian brain, OTRs and V1aRs are broadly expressed in various and often distinct regions^17,25–29^, while V1bR has a more limited distribution, and V2Rs predominantly function in the peripheral tissues. Studies of brain regions and cell types responsible for mediating OTR or VR signaling in diverse biological contexts have been carried out through pharmacological and cKO analyses. For instance, *OTR* cKO models have identified specific cell types that regulate feeding^30^, social reward^31,32^, social aversive learning^33^, social recognition^34,35^, and fear modulation^36^. However, while OT or OTR signaling is shown to modulate pup-related sensory processing^37,38^ in female mice, the specific neurons and signaling mechanisms responsible for the role of OT in promoting paternal behaviors remain unknown. In addition, due to the structural similarity between AVP and OT (sharing seven of nine amino acid residues), AVP can activate not only its canonical VRs, but also OTRs^25,39^. This potential ligand-receptor crosstalk should be considered when identifying the neural hormones and downstream circuits involved in paternal caregiving behaviors.

In the present study, we first establish the roles of PVH AVP neurons in inhibiting infanticide and promoting caregiving behaviors during the paternal life-stage transition. We then conduct a series of viral-genetic experiments to elucidate functionally the receptors and specific brain regions/neural types responsible for the AVP-induced behavioral changes. Our data reveal the existence of non-canonical AVP-to-OTR crosstalk within specific limbic circuits that plays a critical role in regulating paternal behaviors.

## Results

### PVH AVP neurons suppress pup-directed aggression and can promote paternal caregiving behavior

To examine the functional roles of PVH AVP neurons in paternal behaviors, we performed targeted cell ablation of PVH AVP neurons by injecting a Cre-dependent AAV carrying taCasp3-TEVp^40^ (Fig. 1a). Two weeks after the injection, each *AVP-Cre* male mouse^41^ was crossed with a female mouse and housed together throughout her pregnancy and parturition (Fig. 1b). The taCasp3-injected group showed a significant decrease in *AVP*-expressing neurons, while *OT*-expressing neurons remained unaffected (Fig. 1c and 1d), consistent with the negligible overlap of AVP neurons and OT neurons within the PVH^42,43^. While all control fathers that received the saline injection showed pup retrieval, fathers receiving the taCasp3-encoding AAV displayed not only a reduced ratio of retrieval, but also pup-directed aggression (Fig. 1e–1h). The parental care duration, defined as the duration of animals undergoing either grooming, crouching, or retrieving, was significantly lower in the taCasp3-injected fathers (Fig. 1f). We also defined a parental score, where positive values indicated caregiving behaviors and negative values indicated aggression (see Methods). Consistent with the parental care duration, the parental score was significantly lower in taCasp3-injected fathers compared to the saline-injected fathers (Fig. 1g). Of note, ablating PVH AVP neurons resulted in a more pronounced phenotype than did ablating PVH OT neurons, where the latter mainly ignored pups without showing pup-directed aggression (Supplementary Fig. 1). These results indicate that, similar to PVH OT neurons^19^, PVH AVP neurons are required for paternal caregiving behaviors in father mice.

**Fig. 1.**
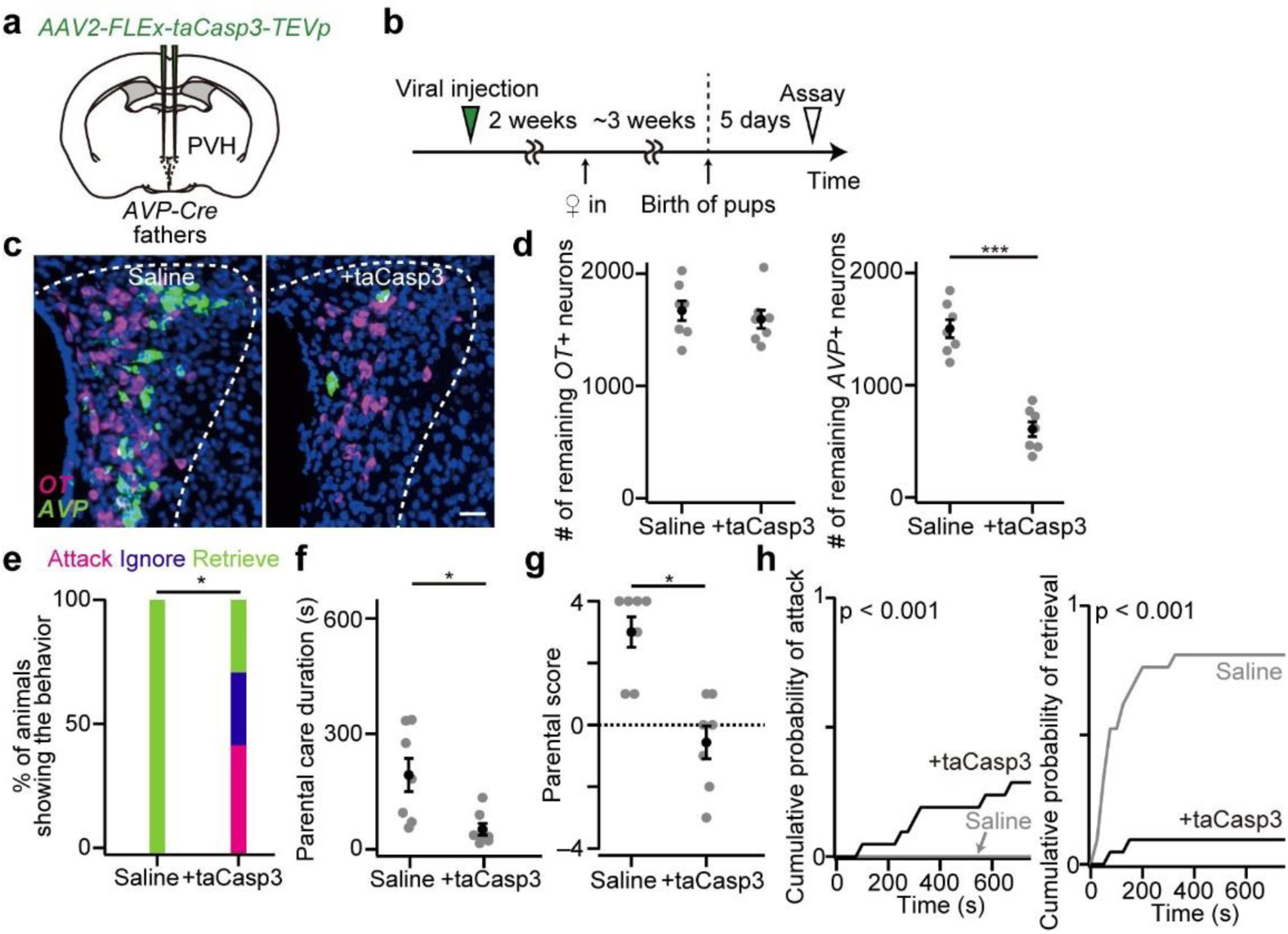
PVH AVP neurons are necessary for suppressing pup-directed aggression in fathers. (a) Schematic of the virus injection. *AAV2-FLEx-taCasp3-TEVp* was injected into the bilateral PVH of *AVP-Cre* mice. (b) Schematic of the time line of the experiment. (c) Representative coronal sections of the PVH without (left) or with (right) *AAV-FLEx-taCasp3-TEVp* injection. *OT* and *AVP in situ* staining are shown in magenta and green, respectively. Blue, DAPI. Scale bar, 30 μm. (d) Number of remaining *OT+* or *AVP+* neurons (***p < 0.001, two-tailed Welch’s *t*-test. n = 7 mice each). (e) Percentage of fathers showing attack, ignore, or retrieve (*p < 0.05, two-tailed Fisher’s exact test). (f) Parental care duration (*p < 0.05, two-tailed Welch’s *t*-test). (g) Parental score (*p < 0.05, two-tailed Mann–Whitney *U*-test). (h) Cumulative probability of pup-directed aggression and pup retrieval. The p-value is shown in the panel (Kolmogorov–Smirnov test). Error bars, SEM. See Supplementary Fig. 1 for more data.

We next explored whether the activation of PVH AVP neurons could suppress infanticide in virgin males, who typically exhibit aggression toward pups. To this end, we chemogenetically activated PVH AVP neurons by expressing *hM3Dq-mCherry* in virgin males (Fig. 2a). Clozapine N-oxide (CNO) or saline as control was administered via intraperitoneal (ip) injection 30 min before the behavioral assay (Fig. 2b). We confirmed comparable expression of hM3Dq in both groups (Fig. 2c). While the saline-injected virgin males showed aggression toward pups, CNO injection significantly inhibited pup-directed aggression and facilitated pup retrieval (Fig. 2d–2g). CNO injection increased the expression of *c-fos* mRNA, a proxy of neural activation, in *Calcr*-expressing (*Calcr+*) neurons in the medial part of the medial preoptic nucleus (MPNm), a crucial center for parental behaviors^13,14^ (Supplementary Fig. 2a–2c). Conversely, neural activity of *urocortin 3*-expressing (*Ucn3+*) neurons in the perifornical area (PeFA), a cell type associated with infanticide^9^, was suppressed (Supplementary Fig. 2d). These findings indicated that PVH AVP neurons suppress pup-directed aggression and facilitate paternal caregiving behaviors by modulating the activity of hypothalamic neural populations associated with parental and infanticidal behaviors.

**Fig. 2.**
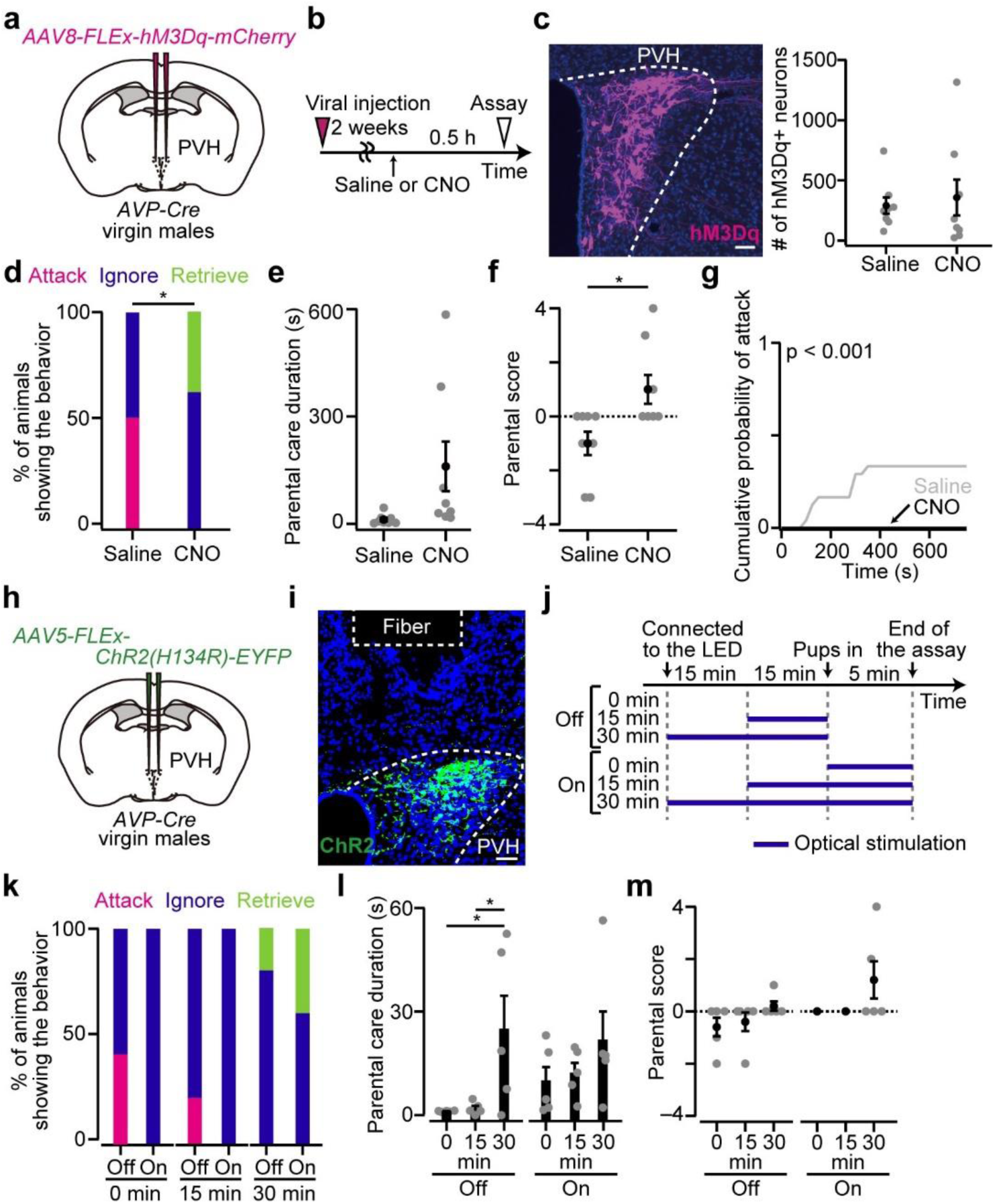
Activation of PVH AVP neurons suppresses pup-directed aggression and can promote caregiving behaviors in virgin males. (a) Schematic of the virus injection. *AAV8-FLEx-hM3Dq-mCherry* was injected into the bilateral PVH of *AVP-Cre* virgin male mice. (b) Schematic of the time line of the experiment. (c) Left, representative coronal section of the PVH. Magenta, hM3Dq-mCherry, blue, DAPI. Scale bar, 50 μm. Right, the number of hM3Dq+ neurons. n = 8 mice each. (d) Percentage of virgin males showing attack, ignore, or retrieve (*p < 0.05, two-tailed Fisher’s exact test). (e) The parental care duration of CNO-injected mice was longer than that of saline-injected males, but did not reach the level of statistical significance (p = 0.086, two-tailed Welch’s *t*-test). (f) Parental score (*p < 0.05, two-tailed Mann–Whitney *U*-test). (g) Cumulative probability of pup-directed aggression. The p-value is shown in the panel (Kolmogorov–Smirnov test). (h) Schematic of the virus injection. *AAV5-FLEx-ChR2(H134R)-eYFP* was injected into the bilateral PVH of *AVP-Cre* virgin male mice. An optical fiber was further inserted into the PVH. (i) Representative coronal brain section showing the location of the optical fiber and expression of ChR2-eYFP (green) in the PVH. Blue, DAPI. Scale bar, 50 μm. (j) Schematic of the time line of the experiment. Blue bar, blue LED stimulation pulsed at 10 Hz. (k) Percentage of virgin males showing attack, ignore, or retrieve. n = 5 each. (l) Parental care duration (*p < 0.05, one-way ANOVA with post hoc Tukey’s HSD). 30-min-illumination evoked a longer parental care duration, although statistical significance was only found in the Off condition. (m) Parental score. Error bars, SEM. See Supplementary Fig. 2 for more data.

Reduction in the number of PVH AVP neurons reinstated pup-directed aggression in fathers (Fig. 1e), whereas chemogenetic activation of PVH AVP neurons not only suppressed pup-directed aggression, but also evoked retrieval in virgin males (Fig. 2d). Additional promotion of retrieval behavior following chemogenetic activation in virgin males may be attributed to the substantial activation of neural circuits governing the suppression of pup-directed aggression, which concurrently facilitates caregiving behavior^5^. To investigate the effects of PVH AVP neurons on caregiving behaviors with higher temporal resolution, we expressed channelrhodopsin-2 (ChR2) in PVH AVP neurons to activate these neurons optically (Fig. 2h and 2i). Each mouse received stimulation for 0, 15, or 30 min before interacting with pups (Fig. 2j). During the interaction, blue light was applied in the On condition, whereas no stimulation was applied in the Off condition (Fig. 2j). We found that even without prior activation of PVH AVP neurons (0 min), pup-directed aggression was completely suppressed in the On condition (Fig. 2k–2m), indicating that acute activation of PVH AVP neurons is sufficient to inhibit pup-directed aggression. In the 0-and 15-min illumination groups, virgin males in the Off condition often exhibited pup-directed aggression, while those illuminated for a longer duration (30 min) became non-infanticidal and even showed parental behaviors, irrespective of the light condition during the behavioral test (Fig. 2k–2m). No effects on paternal behaviors were found in the eYFP control groups (Supplementary Fig. 2e–2h). Together, these results indicate that PVH AVP neurons can modulate paternal behaviors in a scalable manner, depending on the duration of their activation.

### Role of OTRs in mediating AVP neuron-induced paternal behaviors

Having established the functional role of PVH AVP neurons in paternal behaviors, we next examined the downstream receptors responsible for this effect through pharmacological manipulation. We administered an OTR or V1aR antagonist via icv injection while activating PVH AVP neurons with hM3Dq (Fig. 3a–3c). Each experimental group showed a similar number of hM3Dq+ neurons (Fig. 3d). In the absence of AVP neuron activation (ip injection of saline), neither OTR antagonist nor V1aR antagonist administration resulted in substantial changes in behavioral phenotypes (Fig. 3e–3h). Pup-directed aggression was predominant across these groups, with V1aR antagonist application slightly enhancing pup-directed aggression (Fig. 3g and 3h). Consistent with Fig. 2, virgin males receiving CNO did not display pup-directed aggression after receiving an icv injection of saline (Fig. 3e–3h). The icv application of V1aR antagonist significantly resumed pup-directed aggression and reduced parental care duration (Fig. 3e–3h), suggesting that AVP neuron-induced paternal behaviors are partly mediated by V1aRs. However, the application of OTR antagonist reinstated pup-directed aggression more intensively compared with V1aR antagonist (Fig. 3f and 3g), suggesting that PVH AVP neuron-induced paternal behaviors involve OTRs.

**Fig. 3.**
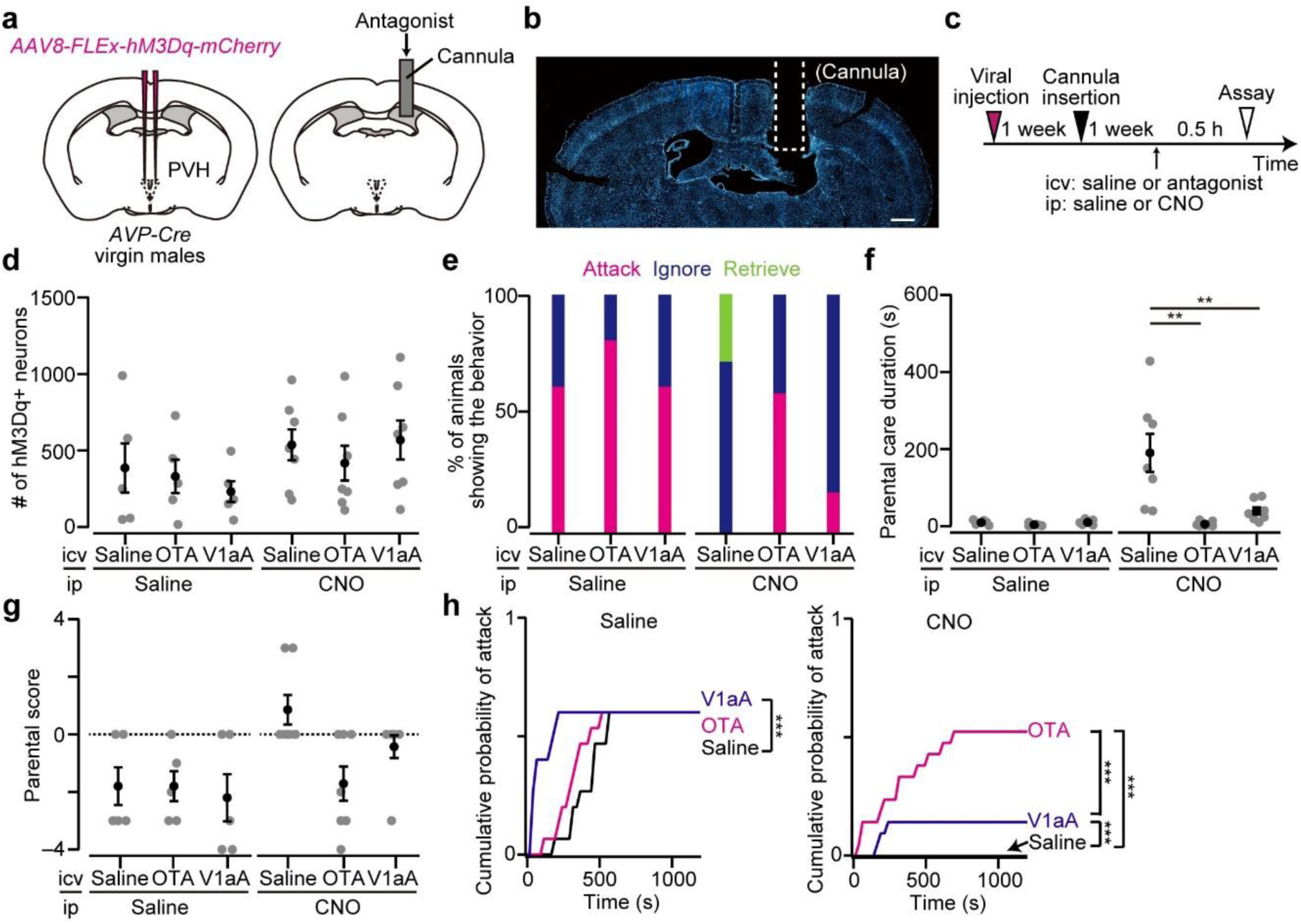
PVH AVP neuron-induced paternal behaviors are suppressed by an OTR antagonist. (a) Schematic of the experiment. *AAV-FLEx-hM3Dq-mCherry* was injected into the bilateral PVH of *AVP-Cre* virgin male mice. OTR antagonist (OTA) or V1aR antagonist (V1aA) was injected into the ventricle through the cannula. (b) Representative coronal section showing the position of the cannula. Blue, DAPI. Scale bar, 500 μm. (c) Schematic of the time line of the experiment. Icv injection of OTA or V1aA was performed under isoflurane anesthesia. Note that the icv injection took approximately 3 min (Methods). (d) The number of neurons expressing hM3Dq was not statistically different (p > 0.46, one-way ANOVA). n = 5 each for ip injection of saline, n = 7 each for ip injection of CNO. (e) Percentage of animals showing attack, ignore, or retrieve. (f) Parental care duration (**p < 0.01, one-way ANOVA with post hoc Tukey’s HSD). (g) Parental score was not significantly different (saline, p > 0.79, CNO, p > 0.06, Kruskal-Wallis test). (h) Cumulative probability of pup-directed aggression (***p < 0.001, Kolmogorov–Smirnov test with Bonferroni correction). Error bars, SEM. See Supplementary Figs. 2 and 3 for more data.

We conducted similar experiments following the chemogenetic activation of PVH OT neurons (Supplementary Fig. 2i–2n). PVH OT neuron-induced facilitation of paternal caregiving behaviors remained unaffected by icv administration of V1aR antagonist, whereas application of OTR antagonist abolished these effects. Thus, paternal behaviors induced by chemogenetic activation of both PVH AVP and OT neurons may be mediated by downstream OTRs somewhere in the brain.

Anatomically, OT neurons receive monosynaptic inputs from AVP neurons within the PVH^19^. Given that OT neurons in the PVH facilitate parental behaviors in male mice^19^, these findings may be explained by the AVP-mediated activation of PVH OT neurons, subsequently activating OTRs. However, the following observations challenge this scenario. First, chemogenetic activation of PVH AVP neurons did not impact the activity of PVH OT neurons as assessed by fiber photometry-based Ca^2+^ imaging (Supplementary Fig. 3a–3f).

We used tail pinch^44^ as a method to assess the recording quality (Supplementary Fig. 3d). These data suggest that, despite the existence of anatomical connections from AVP neurons to OT neurons within the PVH, under our experimental conditions, the functional impacts of this connection are minimal. Second, we found that icv injection of AVP into *OT* KO (*OT^−/−^*) virgin males^19^ reduced the pup-directed aggression to a level similar to that observed in control *OT^+/+^* virgin males (Supplementary Fig. 3g–3l), suggesting that AVP does not require OT release to inhibit pup-directed aggression. Taken together, our data support a scenario wherein AVP neuron-induced paternal caregiving behaviors are mediated through non-canonical AVP-to-OTR crosstalk signaling.

### OTRs in the preoptic area (POA) mediate AVP and OT neuron-induced paternal behaviors

To substantiate our pharmacological data and identify the specific brain regions where OTRs mediate AVP neuron-induced paternal caregiving behaviors, we performed region-specific cKO of the *OTR* gene by injecting *AAV-Cre* into *OTR flox/flox* mice^45^ while chemogenetically activating PVH AVP neurons. Given the known role of the hypothalamus–POA circuit in parental behaviors^3,5^ and its abundant expression of OTRs^13,46,47^, we focused on the OTR neurons in these areas. We first used serotype 9 of *AAV-Cre*, which enables relatively broad *OTR* cKO^30^. In saline-injected control mice without *OTR* cKO, chemogenetic activation of PVH AVP neurons suppressed pup-directed aggression in virgin males (Supplementary Fig. 4). Specifically targeting *OTR* cKO in the “posterior” hypothalamus, including the dorsomedial hypothalamus, lateral hypothalamic area, and ventromedial hypothalamus, resulted in minimal effects on AVP neuron-induced paternal behaviors (Supplementary Fig. 4a–4g). In sharp contrast, *OTR* cKO in the POA disrupted paternal caregiving behaviors and reinstated pup-directed aggression (Supplementary Fig. 4c–4g). These data i) underscore the essential role of OTRs in mediating AVP neuron-induced paternal behaviors and ii) highlight the presence of responsible OTR neurons within the POA.

The POA contains multiple nuclei, and OTR neurons are widely distributed within these regions^13,46,47^. To narrow down which OTR neurons are responsible for mediating AVP neuron-induced paternal behaviors, we utilized serotype 2 of *AAV-Cre*, which enables precise spatial control over cKO at the single nucleus level^30^ (Fig. 4a). Specifically, we focused on the MPNm as a center for parental behaviors^3^ and the anterior commissural nucleus (ACN), a POA subregion activated in virgin males and fathers showing paternal behaviors^7^. We visualized *OTR* expression by RNAscope and defined cells with three or more *OTR* RNAscope dots as *OTR*-expressing (*OTR+*) cells^30^ (Methods; Fig. 4b and 4c). Each group showed comparable hM3Dq expression (Fig. 4d), with a significant reduction in *OTR* expression upon *Cre* expression (Fig. 4e). Virgin males with *OTR* cKO restricted to the MPNm or ACN displayed a lower ratio of AVP neuron-induced pup retrieval (Fig. 4f). The effects were slightly more pronounced with ACN-specific *OTR* cKO, resulting in a significant decrease in parental care duration and an increase in pup-directed aggression (Fig. 4f–4i). However, the *OTR* cKO restricted to the MPNm or ACN only partially impaired AVP neuron-induced paternal behaviors compared with pan-POA cKO (Supplementary Fig. 4c– 4g), suggesting that multiple OTR neurons within the POA synergistically mediate the shift in paternal behavior.

**Fig. 4.**
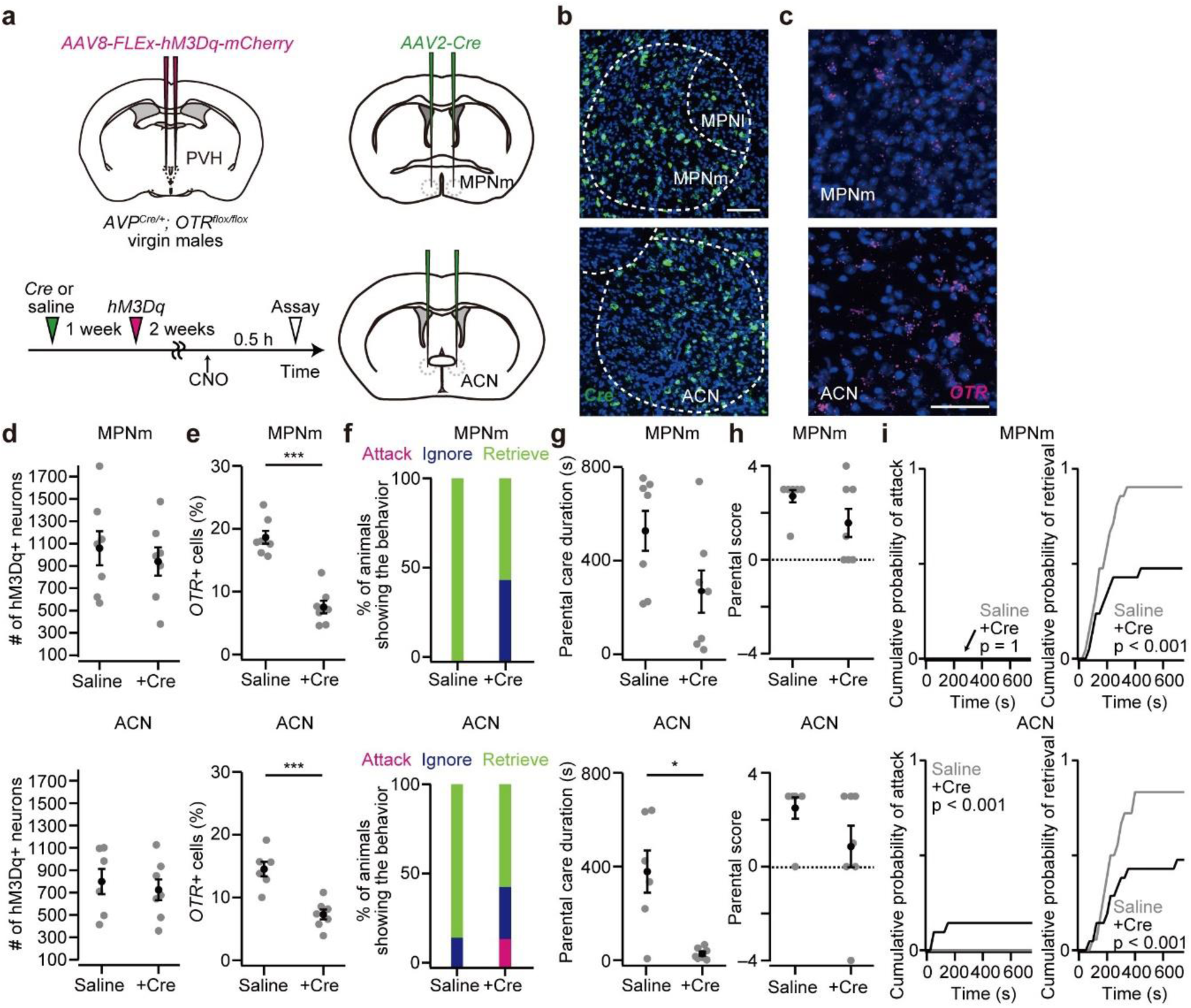
*OTR* in the POA is required for AVP neuron-induced parental behavior. (a) Schematic of the virus injection. *AVP-Cre/+; OTR flox/flox* virgin males were used for the experiment. *AAV8-FLEx-hM3Dq-mCherry* was injected into the bilateral PVH to activate AVP neurons chemogenetically. *AAV2-Cre* was further injected into the bilateral MPNm or ACN to perform cKO of *OTR*. The AAVs were injected as depicted on the time line. (b) Representative coronal sections of the MPNm (top) or ACN (bottom). *Cre in situ* staining is shown in green. Blue, DAPI. MPNl, lateral part of the medial preoptic nucleus. Scale bar, 50 μm. (c) Representative coronal sections showing the MPNm and ACN from a saline-injected mouse. *OTR* mRNA was visualized by RNAscope (magenta). Blue, DAPI. Scale bar, 30 μm. (d) Number of neurons expressing hM3Dq in the PVH. Mice that had 100 or more hM3Dq+ neurons were used for this experiment as they show a higher ratio of retrieval. Top, those mice received *AAV-Cre* injection into the MPNm; bottom, into the ACN. MPNm, n = 7 mice each; ACN, n = 6 and 7 mice for saline and +Cre, respectively. (e) Fraction of DAPI+ cells expressing *OTR* (***p < 0.001, two-tailed Welch’s *t*-test). (f) Percentage of virgin males showing attack, ignore, or retrieve. Of note, although *OTR* in AVP neurons might be deleted in this experimental condition, CNO injection consistently facilitated paternal behaviors in mice that received saline injection into the MPNm or ACN. (g) Parental care duration (*p < 0.05, two-tailed Welch’s *t*-test). (h) Parental score. (i) Cumulative probability of pup-directed aggression and pup retrieval. The p-value is shown in the panel (Kolmogorov–Smirnov test). Error bars, SEM. See Supplementary Figs. 4 and 5 for more data.

Similar experiments were conducted following the chemogenetic activation of PVH OT neurons (Supplementary Fig. 5). *OTR* cKO restricted to the MPNm or ACN (Supplementary Fig. 5a–5g) disrupted PVH OT neuron-induced pup retrieval in virgin males while slightly reinstating pup-directed aggression. Nevertheless, the paternal behaviors induced by PVH OT neurons were not entirely suppressed by these targeted *OTR* cKOs, suggesting OTR signal redundancy. Overall, our data identify OTRs within the MPNm and ACN as key players involved in paternal behaviors triggered by AVP and OT neurons in the PVH, suggesting convergence of these two hormonal signals at downstream targets within the POA.

PVH OT and AVP neurons project their axons to various brain regions, including the POA^35,48^. Having established the functional links from PVH AVP or OT neurons to POA OTR neurons, we aimed to examine their anatomical connections. We generated starter cells for rabies virus-based retrograde transsynaptic tracing^49^ targeting *OTR*+ neurons in the MPNm or ACN using *OTR-iCre* mice^50^. Within the PVH, we found rabies-GFP-positive presynaptic neurons that overlapped with those expressing *OT* or *AVP* mRNA (Supplementary Fig. 6a– 6c). Among all presynaptic neurons in the PVH, the proportion of *OT+* and *AVP+* neurons was similar; approximately 10% and 20% of input neurons to MPNm and ACN *OTR*+ starter cells, respectively, were OT or AVP neurons (Supplementary Fig. 6d). These results suggest that *OTR+* neurons in the MPNm and ACN receive direct inputs from both OT and AVP neurons in the PVH. Of note, *OTR+* neurons in the ACN predominantly consisted of *vGAT*-expressing inhibitory neurons, partly co-expressing *somatostatin* (*SST*), *ets variant 1* (*Etv1*), and *tachykinin 2* (*Tac2*)^13^ (Supplementary Fig. 6e–6i). These results provide insights into the anatomical and cellular basis of connectivity from the PVH to POA, which underlie the facilitation of paternal behaviors in male mice.

### MPNm and ACN OTR neurons mediate paternal caregiving behaviors

If OTR neurons in the MPNm or ACN integrate inputs from PVH OT and AVP neurons, activating these neurons alone in virgin males using chemogenetic methods, without manipulation of the PVH, is sufficient to enhance paternal caregiving behaviors. To test this idea, we used *OTR-iCre* mice^50^ to express *hM3Dq* (Fig. 5a–5d). Activation of *OTR*+ neurons in either the MPNm or ACN effectively suppressed pup-directed aggression and promoted parental caregiving behaviors in virgin males (Fig. 5e–5h). Stimulating ACN *OTR+* neurons showed a slightly stronger effect, leading to a higher ratio of pup retrieval and longer durations of parental care (Fig. 5e and 5f). These results demonstrate the capability of ACN and MPNm OTR neurons to promote paternal caregiving behaviors.

**Fig. 5.**
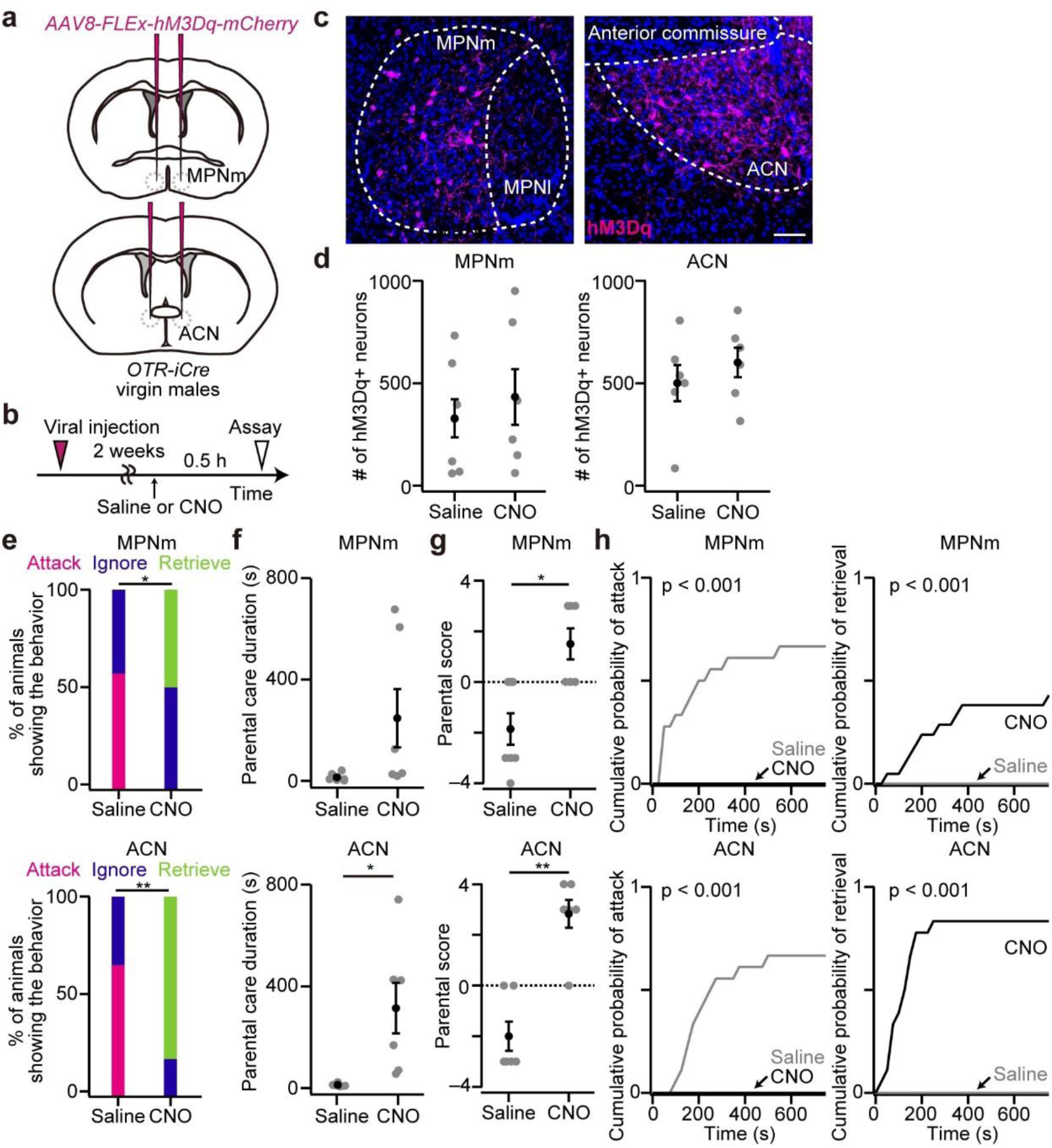
Chemogenetic activation of MPNm or ACN OTR neurons facilitates parental behaviors in virgin males. (a) Schematic of the virus injection. *AAV8-FLEx-hM3Dq-mCherry* was injected into the bilateral MPNm or ACN of *OTR-iCre* virgin males. (b) Schematic of the time line of the experiment. (c) Representative coronal sections of the MPNm (left) or ACN (right). Magenta, hM3Dq-mCherry, Blue, DAPI. MPNl, lateral part of the medial preoptic nucleus. Scale bar, 50 μm. (d) Number of hM3Dq+ neurons. MPNm, n = 7 and 6 for saline and CNO, respectively; ACN, n = 6 each. (e) Percentage of males showing attack, ignore, or retrieve (*p < 0.05, **p < 0.01, two-tailed Fisher’s exact test). (f) Parental care duration (*p < 0.05, two-tailed Welch’s *t*-test). (g) Parental score (*p < 0.05, **p < 0.01, two-tailed Mann–Whitney *U*-test). (h) Cumulative probability of pup-directed aggression and pup retrieval. The p-value is shown in the panel (Kolmogorov–Smirnov test). Error bars, SEM. See Supplementary Fig. 6 for more data.

Lastly, we aimed to investigate the native roles of OTRs in the POA in regulating paternal behavioral changes under physiological conditions. To accomplish this, we assessed the functional contribution of OTRs in the MPNm or ACN to parental behaviors by cKO of the *OTR* gene from these regions of male mice (Fig. 6a and 6b). Injection of serotype 2 of *AAV-Cre* into both the MPNm and ACN significantly reduced the number of *OTR+* cells in these regions in fathers (Fig. 6c–6e). Compared with saline-injected control fathers, who all exhibited paternal caregiving behaviors, *OTR* cKO fathers ignored pups and displayed significantly reduced durations of parental care (Fig. 6f–6h). Notably, targeted *OTR* cKO in either the MPNm or ACN resulted in milder phonotypes, with *OTR* cKO in the ACN leading to a more pronounced impairment of paternal behaviors (Fig. 6i–6m). Thus, OTRs in the POA, particularly in the ACN, are required to promote paternal caregiving behaviors.

**Fig. 6.**
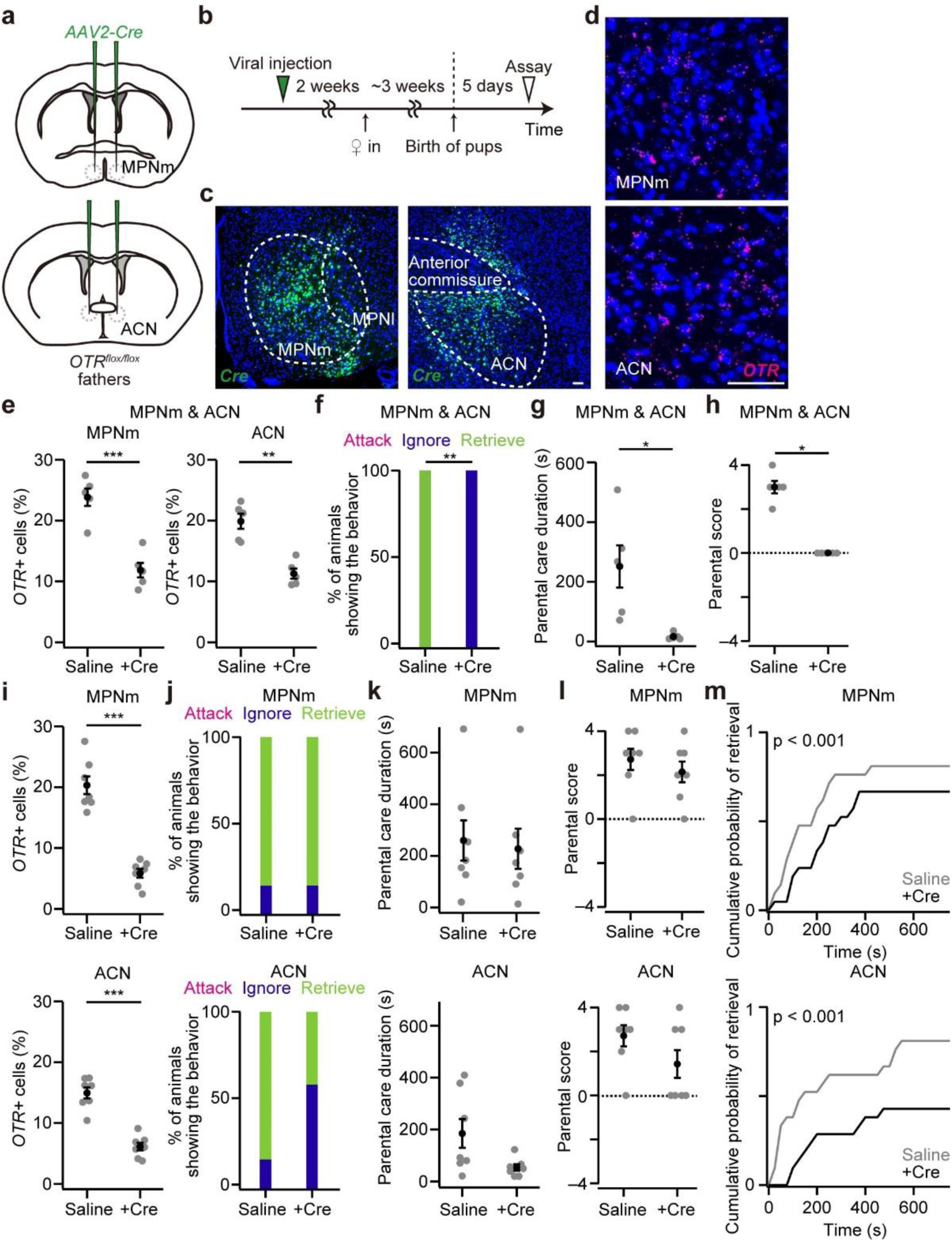
*OTR* in the POA is required for paternal caregiving behaviors. (a) Schematic of the virus injection. *AAV-Cre* was injected into the bilateral MPNm and/or ACN of father mice. (b) Schematic of the time line of the experiment. (c) Representative coronal sections of the MPNm (left) or ACN (right). *Cre in situ* staining is shown in green. Blue, DAPI. MPNl, lateral part of the medial preoptic nucleus. Scale bar, 50 μm. (d) Representative coronal sections showing the MPNm and ACN from a saline-injected mouse. *OTR* mRNA was visualized by RNAscope (magenta). Blue, DAPI. Scale bar, 50 μm. (e) Fraction of DAPI+ cells expressing *OTR* in mice that received *AAV-Cre* injection into both the MPNm and ACN (***p < 0.001, two-tailed Welch’s *t*-test. n = 5 mice each). (f) Percentage of fathers showing attack, ignore, or retrieve (**p < 0.01, two-tailed Fisher’s exact test). (g) Parental care duration (*p < 0.05, two-tailed Welch’s *t*-test). (h) Parental score (*p < 0.05, two-tailed Mann–Whitney *U*-test). (i) Fraction of DAPI+ cells expressing *OTR* in mice that received *AAV-Cre* injection into the MPNm (top) or ACN (bottom) (***p < 0.001, two-tailed Welch’s *t*-test. n = 7 mice each). (j) Behavioral performance of fathers that received *AAV-Cre* injection into the MPNm (top) or ACN (bottom). n = 7 mice each. (k) Parental care duration. (l) Parental score. (m) Cumulative probability of pup retrieval. The p-value is shown in the panel (Kolmogorov– Smirnov test). Error bars, SEM. See Supplementary Fig. 6 for more data.

## Discussion

Despite a growing body of research investigating the neural mechanisms underlying paternal caregiving behaviors^5^, the precise processes that drive the transition from infanticide to caregiving upon fatherhood remain poorly understood. This study revealed several key findings: 1) PVH AVP neurons suppress infanticide and are capable of promoting caregiving behaviors; 2) a substantial portion of AVP neuron-induced paternal behaviors is mediated by OTRs in the POA, specifically in the MPNm and ACN; and 3) these OTR neurons play a critical role in regulating paternal caregiving behaviors. Here, we discuss the biological insights yielded by our study, along with its limitations.

While classical studies in female rats have suggested the involvement of AVP in maternal behaviors^23,51^, the function of AVP neurons in male caregiving behaviors has remained unclear. Our cell-type-selective ablation experiments revealed that PVH AVP neurons are essential for suppressing infanticide, whereas PVH OT neurons are needed for paternal caregiving behaviors (Fig. 1 and Supplementary Fig. 1). Given that paternal behavioral transition occurs in a stepwise manner, from infanticidal to ignoring (Step I) and then to parenting (Step II)^19^, AVP neurons predominately influence Step I, whereas OT neurons become prominent in Step II. Under physiological conditions, both AVP and OT neurons are required for the distinct aspects of paternal behavioral change, suggesting parallel hormonal systems. However, our gain-of-function experiments (Fig. 2 in this study and Fig. 2 in ref. 19) suggest a different perspective: activating either PVH AVP or OT neurons alone is sufficient to induce a fully paternal state. How do we interpret these complex behavioral observations (Supplementary Fig. 6j)? One plausible explanation is that the activity levels of PVH AVP and OT neurons are scalable: the natural behavioral transition may involve modest activations of both neuron types, which can be decoded by a downstream integrator of these activities. Chemogenetic activation of either PVH AVP or OT neurons may provide a super-threshold activity in this potential integrator.

Our data support a model where POA OTR neurons act as integrators of AVP and OT signaling during paternal transition. First, our neural epistatic analyses demonstrated that POA OTRs are essential for AVP-or OT-induced paternal behaviors (Fig. 4, Supplementary Figs. 4 and 5). Second, the chemogenetic activation of POA OTR neurons, especially in the ACN or MPNm, was sufficient to trigger full paternal behavior in male mice (Fig. 5), mimicking the effects of PVH AVP or OT neuron activation. Third, POA OTR neurons received direct axonal projections from both PVH AVP and OT neurons (Supplementary Fig. 6). Lastly, POA *OTR* cKO resulted in an incomplete execution of paternal caregiving behaviors (Fig. 6). Collectively, these lines of evidence support the idea that the scalable activation of POA OTR neurons acts as an integrator of AVP and OT signaling, thereby facilitating the paternal behavioral transition in male mice (Supplementary Fig. 6k). However, we do not exclude the potential contributions of other systems, such as AVP-to-V1aR signaling (Fig. 4) and OT-to-OTR signaling in the brain regions outside the POA, such as the AHi^10^. These alternative pathways may positively influence paternal behaviors and explain the relatively mild phenotype observed in the POA-specific *OTR* cKO fathers (Fig. 6). Future investigations should aim to elucidate the relative roles of multiple widespread systems in mediating OT or AVP signaling during the paternal transition.

Previous studies have postulated potential crosstalk between OT or AVP and non-canonical receptors. *In vitro* binding assays have consistently suggested that the binding affinity of AVP to OTR is comparable to the canonical OT-to-OTR binding, whereas the affinity of OT to VRs is lower^24,25,39^. Recent structural analyses have started to shed light on these binding properties^52^. Consistently, a pharmacological study reported that OTRs, rather than V1aRs, mediate AVP-induced uterine contraction in female mice^53^. Regarding the pain relief functions of OT, even V1aRs have been shown to mediate OT signaling^54^, despite their lower affinity. Our pharmacological and cKO data expand the scope of AVP-to-OTR crosstalk in regulating the paternal behavioral transition, highlighting the functional localization of this crosstalk signaling in the POA. In addition, our data showing that PVH OT neurons primarily use canonical OTRs, not V1aRs (Supplementary Fig. 2), align with known binding affinities, where OT-to-OTR binding is much stronger than OT-to-V1aR^24,25,39^.

We acknowledge two major limitations of the present study. First, although our study highlights the functional importance of PVH AVP and OT neurons in paternal behaviors, when, how, and to what extent these two hormonal systems are activated naturally during the transition to fatherhood remains unclear. Specifically, while PVH OT neurons are known to be active during mating in male mice^55^ and paternal behaviors in male mandarin voles^18^, the activity dynamics of PVH AVP neurons during social behaviors are unknown and thus warrant future investigation. Similarly, gaining a better understanding of the activity dynamics of POA OTR neurons during paternal transition and paternal caregiving behaviors is crucial. Second, it is unclear how POA OTR neurons, particularly those located in the ACN and MPNm, contribute to paternal behaviors. Given their spatial proximity, these OTR neurons are well-positioned to integrate hormonal signals into the parental behavioral centers, such as galanin/Esr1 neurons^11,56^ and Calcr neurons^13,14^ in the MPNm. Future studies are needed to elucidate the detailed architecture and functional properties of the local circuitry connecting OTR neurons with these parental centers in the male POA.

## Methods

### Animals

All animals were housed under a 12-h light/12-h dark cycle with *ad libitum* access to food and water. Wild-type C57BL/6J mice were purchased from Japan SLC. *OT-Cre* (Jax# 024234) and *AVP-Cre* (Jax# 23530) were purchased from the Jackson Laboratory. The *OTR^flox/flox^* mouse line^45^ was provided by Dr. Katsuhiko Nishimori and the *OTR-iCre* mouse line (RIKEN BRC11687; Jax# 037578)^50^ by Drs. Yukiko U. Inoue and Takayoshi Inoue. The mouse line with whole-body KO of *OT* was described previously^19^. All experimental procedures were approved by the Institutional Animal Care and Use Committee of the RIKEN Kobe branch.

### Viral preparations

We obtained the following AAV vectors from Addgene (titer is shown as genome particles [gp] per ml): AAV serotype 9 *hSyn-Cre* (#105555, 2.3 × 10^13^ gp/ml), AAV serotype 8 *hSyn-FLEx-hM3Dq-mCherry* (#44361, 3.2 × 10^13^ gp/ml), AAV serotype 8 *hSyn-FLEx-mCherry* (#50459, 1.7 × 10^13^ gp/ml), AAV serotype 5 *Ef1a-FLEx-eYFP* (#27056, 1.3 × 10^13^ gp/ml), and AAV serotype 5 *Ef1a-FLEx-hChR2(H134R)-eYFP* (#20298, 2.1 × 10^13^ gp/ml). AAV serotype 2 *Ef1a-FLEx-taCasp3-TEVp* (3.4 × 10^12^ gp/ml)^40^ and AAV serotype 2 *CMV-Cre-GFP* (7.1 × 10^12^ gp/ml) were purchased from the University of North Carolina (UNC) viral core. The AAV serotype 5 *CAG-FLEx-TVA-mCherry* (2.4 × 10^13^ gp/ml) and AAV serotype 8 *CAG-FLEx-RG* (1.0 × 10^12^ gp/ml) were generated by the UNC vector core using plasmids, as previously described^49^. The AAV (serotype 9) that expresses *GCaMP* driven by *OT* promotor (*OTp*) was also described previously^57^.

### Stereotactic injection

To target AAV or rabies virus into a specific nucleus, stereotactic coordinates were defined for each nucleus based on the Allen Mouse Brain Atlas^58^. Mice were anesthetized with 65 mg/kg ketamine (Daiichi Sankyo) and 13 mg/kg xylazine (X1251; Sigma-Aldrich) via ip injection and head-fixed to stereotactic equipment (Narishige). The following coordinates were used (in mm from the bregma for anteroposterior [AP], mediolateral [ML], and dorsoventral [DV]): PVH, AP −0.8, ML 0.2, DV 4.5; MPNm, AP −0.2, ML 0.2, DV 5.2; ACN, AP 0.0, ML 1.0, DV 4.5; POA, AP −0.2, ML 0.2, DV 4.5; “posterior” hypothalamus, AP –1.7, ML 1.0, DV 5.2. The injected volume was 200 nl at a speed of 50 nl/min unless stated otherwise. After the viral injection, the animal was returned to the home cage.

### Parental behavior assay

#### Assay for virgin males

The behavioral assay was conducted as previously described^19^. In brief, 10–12-week-old virgin males were housed individually for 5–7 days. Each cage contained shredded paper on wood chips. The mouse builds its nest with the papers, typically at a corner of the cage. All experiments were performed under dim fluorescent light. Three 5–7-day-old C57BL/6J pups that had not been exposed to any adult males before the experiment were placed in different corners where the testing male had not built its nest. The introduction of pups marked the beginning of the assay. Males were allowed to interact with the pups freely for 15 min for chemogenetic experiments (Figs. 2, 4, and 5, Supplementary Figs. 4 and 5) and 5 min for optogenetic experiments (Fig. 2 and Supplementary Fig. 2). If any sign of pup-directed aggression was observed, the targeted pup was immediately rescued from the cage. For the behavioral assay with chemogenetic activation, AAV that expresses *hM3Dq* was injected 2 weeks beforehand. Each male was tested only once. At 30 min before the assay, CNO (4930; Tocris) dissolved in saline was administered via ip injection to achieve a dose of 1 mg/kg. A saline-only injection was used as a control. Two identical tubes containing either saline or saline with CNO were prepared, and the experimenter who conducted the behavioral assay, immunostaining, and data analysis was blinded to the contents of the tubes. For the behavioral assay with optogenetic activation, *AAV-FLEx-ChR2(H134R)-eYFP* or *AAV-FLEx-eYFP* was injected into the bilateral PVH of *AVP-Cre* mice 3 weeks before the behavioral assay. At 2 weeks after the viral injection, a 400-μm core, N.A. 0.5 optical fiber (R-FOC-BL400C-50NA; RWD) was implanted approximately 500 μm above the PVH. Blue light (465 nm) emitted from an LED (CLED_465; Doric Lenses) was pulsed at 10 Hz. The optical intensity measured at the tip of the fiber was approximately 127 μW (S120VC sensor; Thorlabs). Each male was tested under only one illumination condition. The behavior of the animals was categorized into ‘Attack’, ‘Ignore’, or ‘Retrieve’ based on the criteria previously described^19^. The duration of animals undergoing either grooming, crouching, or retrieving was scored as parental care duration. The parental score was calculated based on a previously described method^6^ with several modifications. First, each mouse was assigned a score of 0. The number of pups retrieved or attacked by the male was added to or subtracted from this score. If the male retrieved or attacked all three pups within 2 min, one point was further added or subtracted, respectively. Consequently, the maximum value of the parental score is 4, and the minimum score is −4.

#### Assay for fathers

The behavioral assay with fathers was conducted using the same procedure as that for virgin males, with several modifications. An individually housed virgin male (10 weeks old) was paired with a female. The next day, the vaginal plug was checked, and only the male– female pairs that successfully formed a plug were used for further experiments. Males were allowed to cohabit with the mated female until 5 days after the birth of pups. The behavioral assay for the father mouse was conducted 5 days after the birth of pups. Females (mothers) and pups were removed from the home cage 6 h before the assay, leaving only the fathers. Unfamiliar pups (pups unrelated to the resident father), prepared as the assay for the virgin males, were used for the assay.

#### Assay for c-fos expression

The behavioral assay for visualizing *c-fos* expression by *in situ* hybridization (ISH) (Supplementary Fig. 2) was conducted with virgin males. Before the assay, each virgin male was individually housed for 7 days in a cage containing a metallic tea strainer. The assay was performed in the dark. On the test day, three pups were packed into the tea strainer, which prevented them from being attacked by virgin males during the assay. Males were allowed to interact with pups in the tea strainer for 20 min. After 20 min, males were sacrificed as described in the Histochemistry section.

#### Assay with icv injection of chemicals

A 22-gauge guide cannula (C313GA/SPC; Plastics One) was inserted into the third ventricle and fixed with dental cement. A dummy cannula (C313DC/SPC) was inserted into the guide cannula and the mouse was returned to the home cage. At 30 min before the behavioral assay, each mouse was moved to a new cage and anesthetized with 3% isoflurane (Narcobit-E; Natsume Seisakusho). Under anesthesia, a dummy cannula was removed and an internal 28-gauge cannula (C313LI/SPC) was inserted into the ventricle through the guide cannula. OTR antagonist ((d(CH_2_)_5_¹, Tyr(Me)², Thr⁴, Orn⁸, des-Gly-NH_2_^9^)-vasotocin; #4031339, Bachem), V1aR antagonist ([β-Mercapto-β,β-cyclopentamethylenepropionyl^1^, O-me-Tyr^2^, Arg^8^]-vasopressin; V2255, Sigma)^59^, or AVP (#2935, Tocris) was dissolved in saline at 500 μM. Next, 10 μl of the solution was injected at a rate of 10 μl/3 min, controlled by a microsyringe pump (MSP-3D; As One). After the injection, the internal cannula was removed and a dummy cannula was inserted. The mouse was then returned to the home cage.

### Fiber photometry

Fiber photometry recordings were performed as described previously^57^. In brief, each *AVP-Cre* male mouse received 200 nl of a 1:1 mixture of *AAV8-FLEx-hM3Dq-mCherry* and AAV (serotype 9) that expresses *GCaMP6s* driven by *OT* promoter^57^ in the bilateral PVH. At 2 weeks after the viral injection, a 400-μm core, N.A. 0.5 optical fiber (R-FOC-BL400C-50NA; RWD) was implanted approximately 100 μm above the PVH. In this experimental design, ip injection of CNO activates PVH AVP neurons, and the possible response of PVH OT neurons can be detected by GCaMP signals. The mice that showed an increase of GCaMP signals in response to the tail pinch, which activates PVH OT neurons^44^, were used for the experiments. The area under the curve (Supplementary Fig. 3f) was calculated from ΔF/F, where F denotes the average fluorescence intensity at 1 hour before stimulation. The reported values in Supplementary Fig. 3f were obtained by dividing the CNO condition by the saline condition.

### Transsynaptic retrograde tracing

The rabies virus used in this study was prepared with viruses, cell lines, and protocols as previously described^19,60^. In brief, RV*d*G-GFP was prepared using the B7GG cell line (a gift from Ed Callaway) and plasmids (*pCAG-B19N*, *pCAG-B19P*, *pCAG-B19L*, *pCAG-B19G*, and *pSADdG-GFP-F2*; gifts from Ed Callaway), and then pseudotyped using BHK-EnvA cells (a gift from Ed Callaway). The titer of RV*d*G-GFP+EnvA used in this study was estimated to be 4 × 10^9^ infectious particles per ml based on serial dilutions of the virus stock followed by infection of the HEK293-TVA800 cell line (a gift from Ed Callaway).

To initiate the transsynaptic tracing using rabies virus, 100 nl of a 1:1 mixture of AAV5 *CAG-FLEx-TVA-mCherry* and AAV8 *CAG-FLEx-RG*^49^ was injected into the unilateral MPNm or ACN of *OTR-iCre* mice at a speed of 30 nl/min. Two weeks later, 200 nl of RV*dG-GFP*+EnvA was injected into the same brain region. At 1 week after the injection of rabies virus, mice were sacrificed and perfused with PBS followed by 4% PFA in PBS. Then, 20-μm coronal sections were collected. Images were acquired with a 10× (N.A. 0.4) objective lens and cells were counted manually using the ImageJ Cell Counter plugin.

### Histochemistry

Mice were anesthetized with isoflurane and perfused with PBS followed by 4% PFA in PBS. The brain was then post-fixed with 4% PFA overnight. Then, 20-μm coronal brain sections were obtained using a cryostat (Leica). Fluorescent ISH was performed as previously described^19,61^. Sections were treated with TSA-plus Cyanine 3 (NEL744001KT; Akoya Biosciences) or TSA-plus biotin (NEL749A001KT; Akoya Biosciences) followed by streptavidin-Alexa Fluor 488 (S32354; Invitrogen). The primers (5’ – 3’) to produce RNA probes were as follows (the first one, forward primer, the second one, reverse primer):

*OT*, 5’-AAGGTCGGTCTGGGCCGGAGA; 5’-TAAGCCAAGCAGGCAGCAAGC

*AVP*, 5’-ACACAGTGCCCACCTATGCT; 5’-CTCTTGGGCAGTTCTGGAAG

*Cre*, 5’-CCAAGAAGAAGAGGAAGGTGTC; 5’-ATCCCCAGAAATGCCAGATTAC

*Galanin*, 5’-GCTCCCACTGGGCATAAATA; 5’-GCTTGAGGAGTTGGCAGAAG

*Calcr* (part 1), 5’-CTGCTCCTAGTGAGCCCAAC; 5’-AGCAAGTGGGTTTCTGCACT

*Calcr* (part 2), 5’-TCCCAGGAGCTGACCATATC; 5’-TAGCAGCAAGCAAGAGGTCA

*Calcr* (part 3), 5’-TTGCCCTTGGGTGCTATCTA; 5’-AGCAGAAGCGTTTCACACAA

*Ucn3*, 5’-TTGCTTCTCGGCTTACCTGT; 5’-AATTCTTGGCCTTGTCGATG

*c-fos* (part 1), 5’-AGCGAGCAACTGAGAAGACTG; 5’-ATCTCCTCTGGGAAGCCAAG

*c-fos* (part 2), 5’-CCAGTCAAGAGCATCAGCAA; 5’-CATTCAGACCACCTCGACAA

*GFP*, 5’-ACGTAAACGGCCACAAGTTC; 5’-CTTGTACAGCTCGTCCATGC

*SST*, 5’-GTGTGCTCCTATGTGGCTGA; 5’-TCAATTTCTAATGCAGGGTCAA

*Etv1*, 5’-TGTGCCTTGCTGTTTCATTC; 5’-CATCCCTCTTTTGATCCGTTA

*Tac2*, 5’-CCCTGCACTCTTGTCTCTGTC; 5’-CTATGGGGTTGAGGCTGTTC

In *Calcr* and *c-fos* ISH, mixtures of parts 1–3 and parts 1 and 2 were used, respectively.

For the immunohistochemistry, the following reagents were used for primary antibodies: anti-GFP (GFP-1010; Aves Labs; 1:500), anti-RFP (5f8; Chromotek; 1:500), and anti-mCherry (AB0040-200; OriGene; 1:500). Signal-positive cells were detected by anti-chicken Alexa Fluor 488 (703-545-155; Jackson Immuno Research; 1:500), anti-rat Cy3 (712-165-153; Jackson Immuno Research; 1:500), and anti-goat 555 (A32816; Invitrogen; 1:500). Fluoromount (K024; Diagnostic BioSystems) was used as a mounting medium. Brain images, except Fig. 3b, were acquired using an Olympus BX53 microscope equipped with a 10× (N.A. 0.4) objective lens. Fig. 3b was obtained using a slide scanner (AxioScan; Zeiss). Cells were counted manually using the ImageJ Cell Counter plugin.

### RNAscope

*OTR* mRNA was visualized using the RNAscope Multiplex Fluorescent Reagent Kit (323110; Advance Cell Diagnostics [ACD]) according to the manufacturer’s instructions. Next, 20-μm coronal brain sections were made using a cryostat. A probe against *OTR* (Mm-OXTR, 412171; ACD) was hybridized in a HybEZ Oven (ACD) for 2 h at 40 °C, followed by a visualization process with TSA-plus Cyanine 3 (NEL744001KT; Akoya Biosciences; 1:1500). Fluoromount (K024; Diagnostic BioSystems) was used as a mounting medium. Images subjected to the analysis were acquired using an Olympus BX53 microscope equipped with a 10× (N.A. 0.4) objective lens. *OTR* expression was observed as a dot-like structure, and cells showing three or more RNAscope dots were defined as *OTR+*^30^.

### Data analysis

All mean values are reported as mean ± SEM. The statistical details of each experiment, including the statistical tests used, the exact value of n, and what n represents, are shown in each figure legend. The p-values are shown in each figure legend or panel; nonsignificant values are not noted.

## Data and materials availability

All data are available in the main paper and supplementary materials. All materials are available through reasonable request to the corresponding authors.

## Code availability statement

No original code was generated in the course of this study. Any additional information required to reanalyze the data reported in this work paper is available from the Lead Contact upon request.

## Author contributions

K.I. and K.M. conceived and designed the project. K.I., M.H., and K.Y. performed the experiments. K.I. and K.Y. analyzed the data. S.I. generated the pseudotyped rabies virus. Y.U.I. and T.I. provided the *OTR-iCre* mice. K.I. and K.M. wrote the paper.

## Acknowledgments

We thank Katsuhiko Nishimori for the provision of the *OTR flox/flox* mouse line. We also thank Teruhiro Okuyama (the University of Tokyo) and the members of the Miyamichi Lab for the critical reading of the manuscript. This work was supported by the RIKEN Special Postdoctoral Researchers Program, a grant from the Kao Foundation for Arts and Sciences, the IDDI Outstanding Basic and Applied Neuroscience Talent Award, and JSPS KAKENHI (19J00403, 19K16303, and 23K14310) to K.I., and JSPS KAKENHI (20K20589 and 21H02587) to K.M.

---

**Supplementary Fig. 1.**
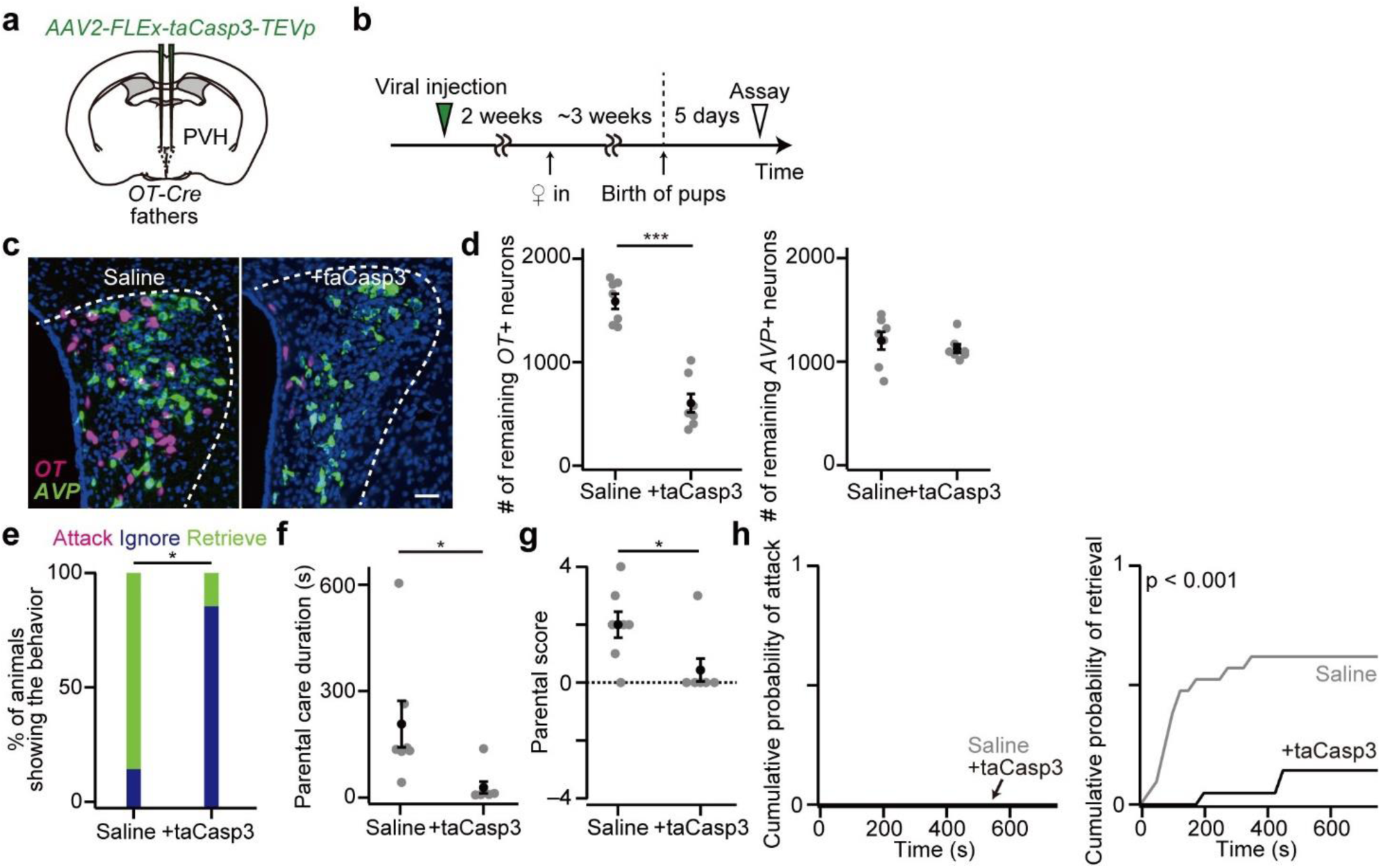
Father mice with cell ablation of PVH OT neurons mostly ignored pups, related to Fig. 1. (a) Schematic of the virus injection. *AAV-FLEx-taCasp3-TEVp* was injected into the bilateral PVH of *OT-Cre* mice. (b) Schematic of the time line of the experiment. (c) Representative coronal sections of the PVH without (left) or with (right) *AAV-FLEx-taCasp3-TEVp* injection. *OT* and *AVP in situ* staining are shown in magenta and green, respectively. Blue, DAPI. Scale bar, 50 μm. (d) Number of remaining *OT+* or *AVP+* neurons (***p < 0.001, two-tailed Welch’s *t*-test. n = 7 mice each). (e) Percentage of fathers showing attack, ignore, or retrieve (*p < 0.05, two-tailed Fisher’s exact test). (f) Parental care duration (*p < 0.05, two-tailed Welch’s *t*-test). (g) Parental score (*p < 0.05, two-tailed Mann–Whitney *U*-test). (h) Cumulative probability of pup-directed aggression and pup retrieval. The p-value is shown in the panel (Kolmogorov–Smirnov test). Of note, we reported similar results in the context of expectant fathers (before the birth of their offspring) in Supplementary Fig. 2 of ref. ^19^. The data presented in this figure are from a new cohort with a behavioral assay conducted 5 days after the birth of pups. Error bars, SEM.

**Supplementary Fig. 2.**
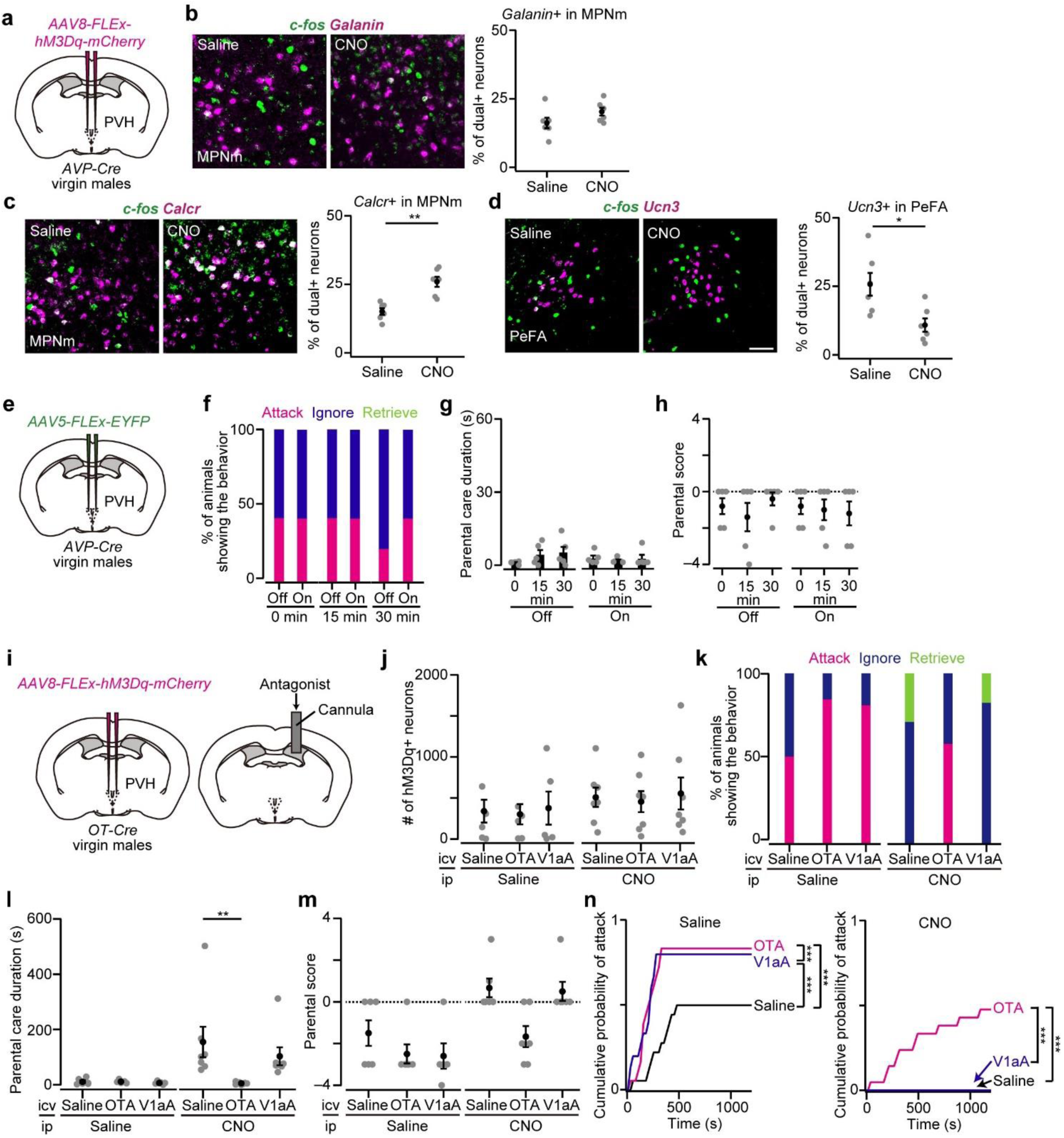
*c-fos* assay with chemogenetic activation of PVH AVP neurons, and more data for optogenetic and icv injection experiments, related to Figs. 2 and 3. (a) Schematic of the virus injection. (b–d) Representative coronal sections of *AVP-Cre* virgin males expressing hM3Dq who interacted with isolated pups in a metal strainer (see Methods). Green represents *c-fos* mRNA. Magenta represents *Galanin* mRNA in the MPNm (b), *Calcr* mRNA in the MPNm (c), or *Ucn3* mRNA in PeFA (d) *in situ* staining (*p < 0.05, **p < 0.01, two-tailed Welch’s *t*-test). n = 6 each. Blue, DAPI. Scale bar, 30 μm. (e) Schematic of the virus injection. *AAV5-FLEx-eYFP* was injected into the bilateral PVH. An optical fiber was further inserted into the PVH. The experimental procedures are the same as those in Fig. 2j. (f) Percentage of virgin males showing attack, ignore, or retrieve. n = 5 mice each. (g) Parental care duration. (h) Parental score. (i) Schematic of the experiment. *AAV-FLEx-hM3Dq-mCherry* was injected into the bilateral PVH of *OT-Cre* mice. OTA, OTR antagonist, V1aA, V1aR antagonist. (j) The number of neurons expressing hM3Dq was not statistically different (p > 0.87, one-way ANOVA). Ip injection of saline, n = 6, 6, and 5 for saline, OTA, and V1aA, respectively. Ip injection of CNO, n = 7 mice each. (k) Percentage of animals showing attack, ignore, or retrieve. (l) Parental care duration (**p < 0.01, one-way ANOVA with post hoc Tukey’s HSD). (m) Parental score was not significantly different (saline, p > 0.46, CNO, p = 0.05, Kruskal-Wallis test). (n) Cumulative probability of pup-directed aggression (***p < 0.001, Kolmogorov–Smirnov test with Bonferroni correction). Error bars, SEM.

**Supplementary Fig. 3.**
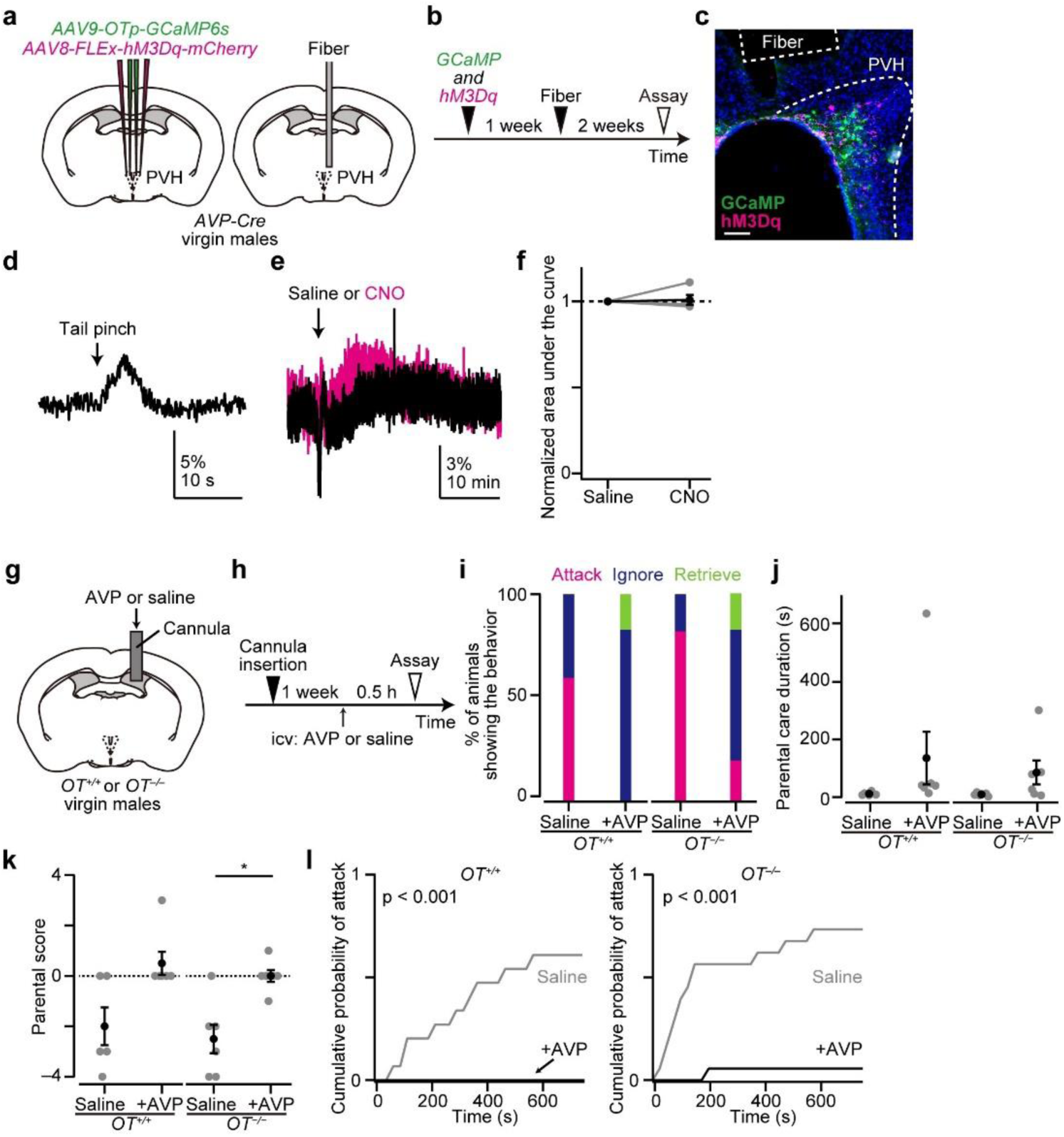
AVP-induced paternal behaviors are independent of OT secretion, related to Figs. 2 and 3. (a) Schematic of the virus injection. AVP neurons express *hM3Dq-mCherry* while OT neurons express GCaMP driven by *OT promotor (OTp)*. (b) Schematic of the time line of the experiment. (c) Representative coronal brain section showing the location of the optical fiber and expression of GCaMP (green) and hM3Dq-mCherry (magenta) in the PVH. Blue, DAPI. Scale bar, 50 μm. (d) Sample recording from OT neurons in response to a tail pinch (arrow). (e) Sample recording from OT neurons with ip injection of saline or CNO. (f) The normalized area under the curve was not affected by CNO application (n = 4). (g) Schematic of the experiment. Saline containing AVP or saline alone was injected into the ventricle of wild-type (*OT^+/+^*) or *OT* KO (*OT^−/−^*) virgin male mice. (h) Schematic of the time line of the experiment. (i) Percentage of animals showing attack, ignore, or retrieve (n = 5 and 6 for saline and AVP in *OT^+/+^*, respectively; n = 6 each in *OT^−/−^*). (j) Parental care duration. (k) Parental score (*p < 0.05, two-tailed Mann–Whitney *U*-test). (l) Cumulative probability of pup-directed aggression. The p-value is shown in the panel (Kolmogorov–Smirnov test). Error bars, SEM.

**Supplementary Fig. 4.**
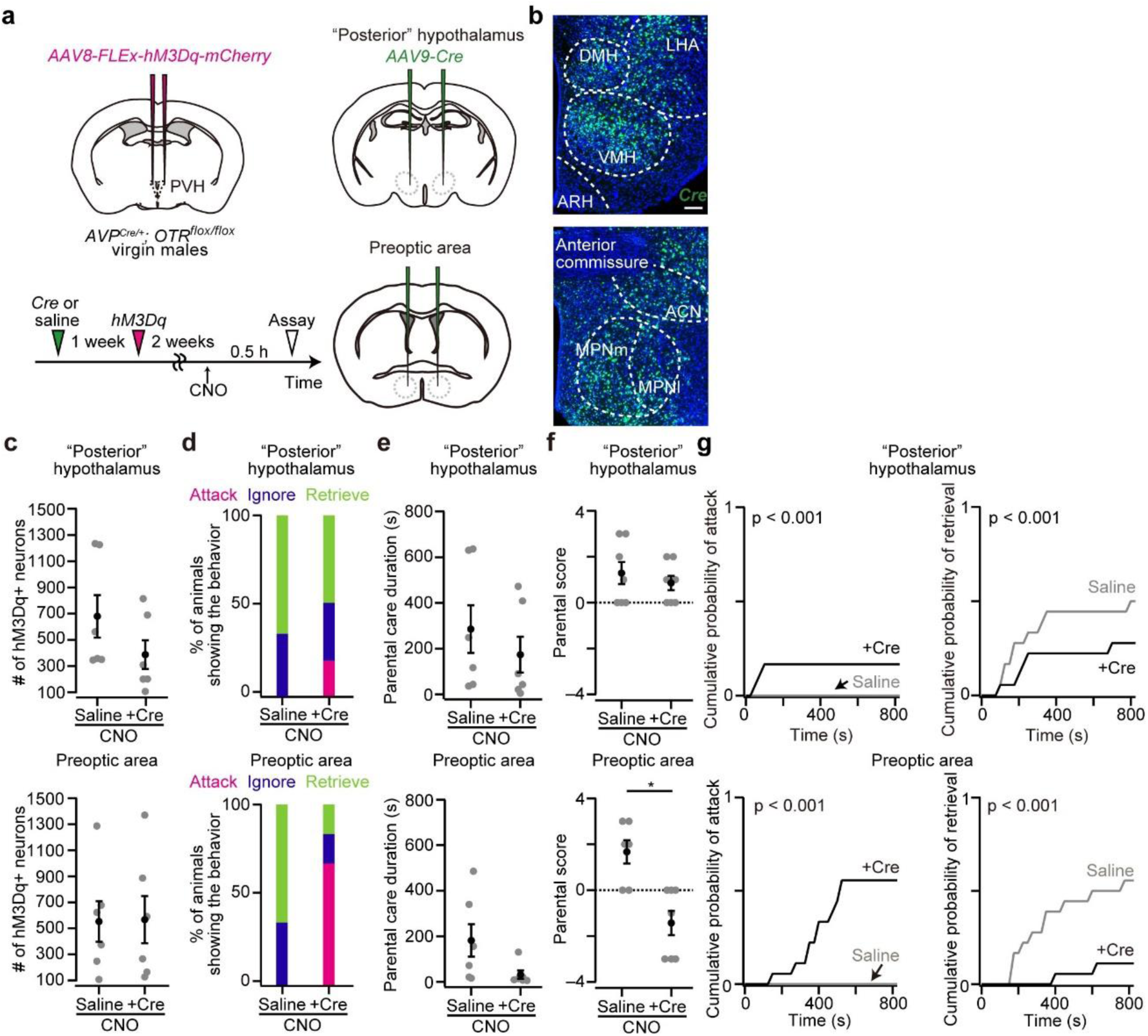
OTR in the POA is required for AVP neuron-mediated paternal behaviors, related to Fig. 4. (a) Schematic of the experiment. *AVP-Cre/+; OTR flox/flox* mice were used for the experiment. *AAV-Cre* was injected into the POA or “posterior” hypothalamus while *AAV-FLEx-hM3Dq-mCherry* was injected into the bilateral PVH. (b) Representative coronal sections. *Cre in situ* staining is shown in green. Blue, DAPI. Scale bar, 50 μm. (c) Number of hM3Dq+ neurons in the PVH (n = 6 each). Mice that had 100 or more hM3Dq+ neurons were used. (d) Percentage of animals showing attack, ignore, or retrieve. (e) Parental care duration. (f) Parental score (*p < 0.05, two-tailed Mann–Whitney *U*-test). (g) Cumulative probability of pup-directed aggression. The p-value is shown in the panel (Kolmogorov–Smirnov test). Error bars, SEM.

**Supplementary Fig. 5.**
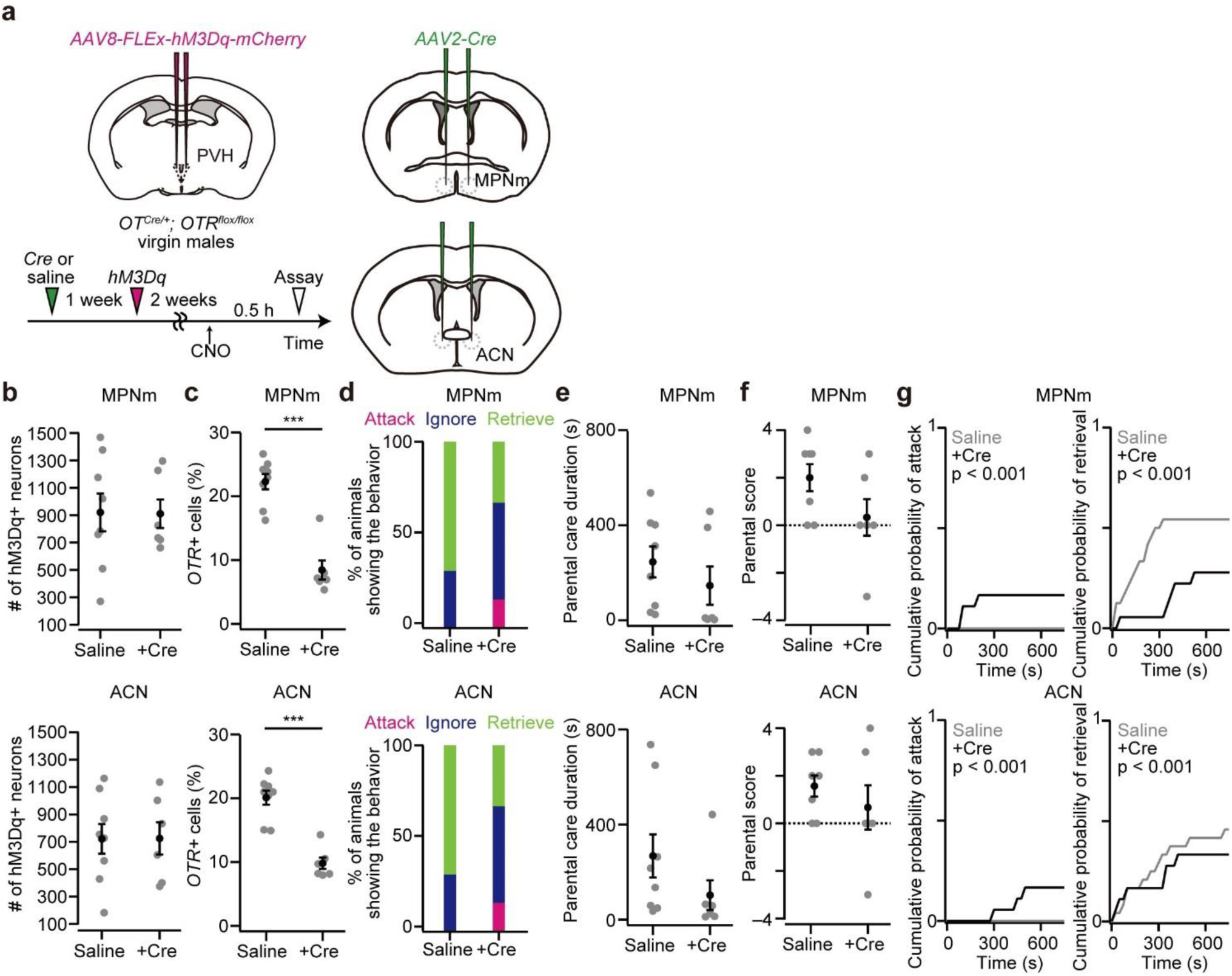
*OTR* in the POA nuclei is partially required for PVH OT neuron-induced paternal behaviors, related to Fig. 4. (a) Schematic of the virus injection and experimental time line. *AAV8-FLEx-hM3Dq-mCherry* was injected into the bilateral PVH, while *AAV2-Cre* was injected into the bilateral MPNm or ACN of *OT-Cre/+; OTR flox/flox* virgin males. (b) Number of neurons expressing hM3Dq in the PVH. Top, data from the mice who received *AAV-Cre* injection into the MPNm; bottom, into the ACN. MPNm, n = 7 and 6 for saline and +Cre, respectively; ACN, n = 7 and 6 for saline and +Cre, respectively. (c) Fraction of DAPI+ cells expressing *OTR* (***p < 0.001, two-tailed Welch’s *t*-test). (d) Percentage of virgin males showing attack, ignore, or retrieve. Of note, *OTR* may be deleted in OT neurons in this experimental condition, which did not affect the induction of pup retrieval upon CNO injection in mice that received saline injection into the MPNm or ACN. (e) Parental care duration. (f) Parental score. (g) Cumulative probability of pup-directed aggression and pup retrieval. The p-value is shown in the panel (Kolmogorov–Smirnov test). Error bars, SEM.

**Supplementary Fig. 6.**
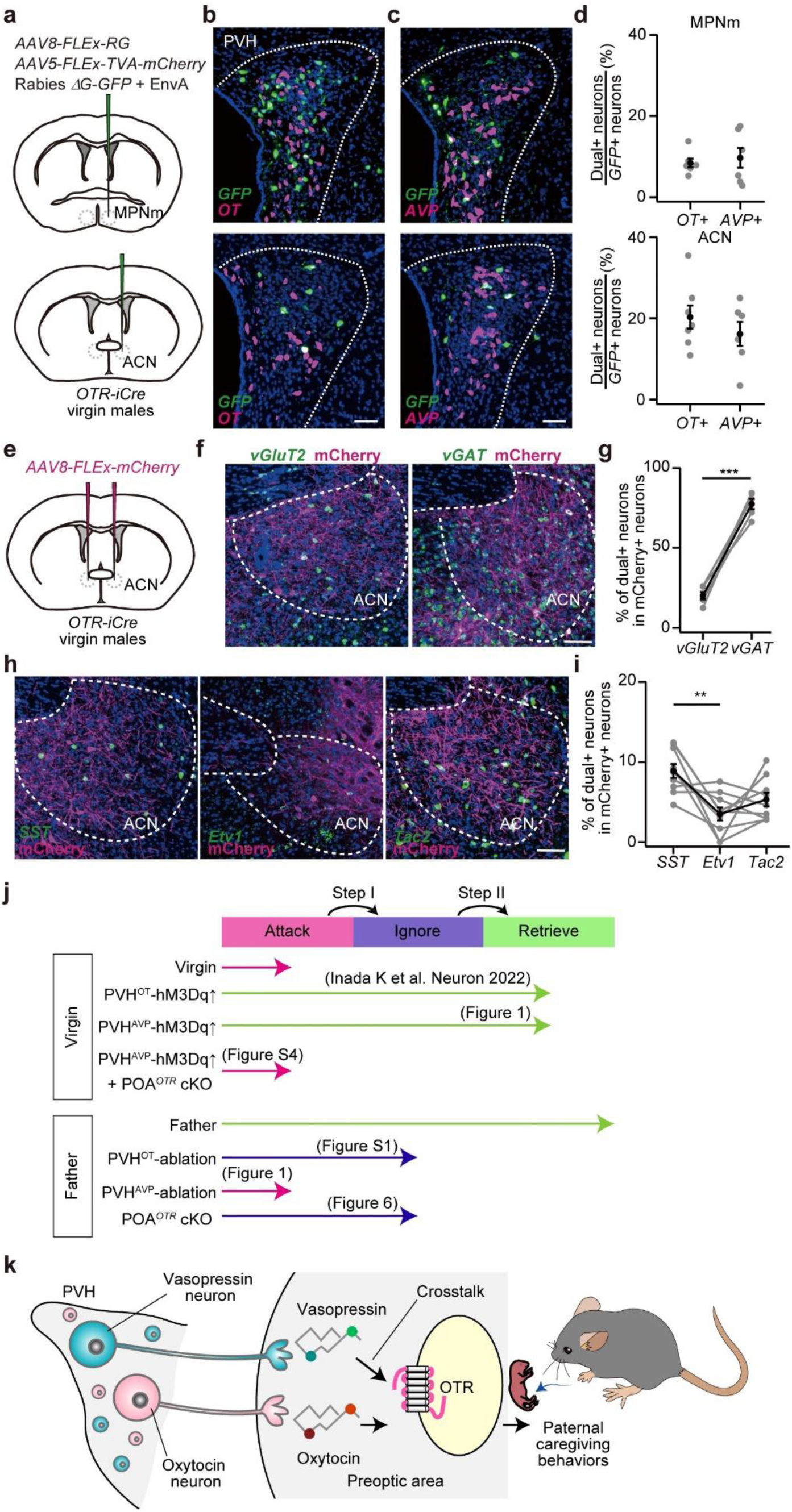
Characteristics of ACN OTR neurons and graphical summary, related to Figs. 5 and 6. (a) Schematic of the virus injection. Presynaptic neurons of *OTR+* neurons in the MPNm (top) or ACN (bottom) are labeled by *GFP* expression. (b, c) Representative coronal sections of the PVH with transsynaptic tracing from the MPNm (top) or ACN (bottom). Green represents *in situ* staining of *GFP*. Magenta represents *in situ* staining of *OT* (b) or *AVP* (c). Blue, DAPI. Scale bar, 30 μm. (d) Percentage of dual-positive neurons among *GFP+* neurons in the PVH. In presynaptic neurons among *OTR+* neurons in the MPNm (top) and *OTR+* neurons in the ACN (bottom), no significant difference was found between *OT+* and *AVP+* (p > 0.93 and p > 0.56 for MPNm and ACN, respectively, Two-tailed Mann–Whitney *U*-test. MPNm, n = 6 each; ACN, n = 7 and 6 for *OT+* and *AVP+*, respectively). (e) Schematic of the virus injection. *AAV8-FLEx-mCherry* was injected into the bilateral ACN of *OTR-iCre* virgin males. (f) Representative coronal sections of the ACN. Green shows *vGluT2* (left) or *vGAT* (right) *in situ* staining. Magenta, mCherry. Blue, DAPI. Scale bar, 50 μm. (g) Fraction of mCherry+ neurons co-expressing *vGluT2* or *vGAT* (n = 5 mice; ***p < 0.001, paired *t*-test). (h) Representative coronal sections of the ACN showing *SST* (left), *Etv1* (middle), or *Tac2* (right) mRNA (green). Magenta, mCherry. Blue, DAPI. Scale bar, 50 μm. (i) Fraction of mCherry+ neurons co-expressing *SST*, *Etv1*, or *Tac2* (n = 9 mice; **p < 0.01, one-way ANOVA with a post hoc paired *t*-test with Bonferroni correction). (j) Summary of behavioral phenotypes. (k) Graphical abstract.

## References

1. Lukas, D., and Huchard, E. (2019). The evolution of infanticide by females in mammals. Philos Trans R Soc Lond B Biol Sci 374, 20180075. 10.1098/rstb.2018.0075.

2. Lukas, D., and Huchard, E. (2014). Sexual conflict. The evolution of infanticide by males in mammalian societies. Science 346, 841–844. 10.1126/science.1257226.

3. Dulac, C., O’Connell, L.A., and Wu, Z. (2014). Neural control of maternal and paternal behaviors. Science 345, 765–770. 10.1126/science.1253291.

4. Elwood, R.W. (1994). Temporal-based kinship recognition: A switch in time saves mine. Behav Processes 33, 15–24. 10.1016/0376-6357(94)90057-4.

5. Inada, K., and Miyamichi, K. (2023). Association between parental behaviors and structural plasticity in the brain of male rodents. Neurosci Res 196, 1–10. 10.1016/j.neures.2023.06.007.

6. Tachikawa, K.S., Yoshihara, Y., and Kuroda, K.O. (2013). Behavioral transition from attack to parenting in male mice: a crucial role of the vomeronasal system. J Neurosci 33, 5120–5126. 10.1523/JNEUROSCI.2364-12.2013.

7. Tsuneoka, Y., Tokita, K., Yoshihara, C., Amano, T., Esposito, G., Huang, A.J., Yu, L.M., Odaka, Y., Shinozuka, K., McHugh, T.J., and Kuroda, K.O. (2015). Distinct preoptic-BST nuclei dissociate paternal and infanticidal behavior in mice. EMBO J 34, 2652–2670. 10.15252/embj.201591942.

8. Chen, P.B., Hu, R.K., Wu, Y.E., Pan, L., Huang, S., Micevych, P.E., and Hong, W. (2019). Sexually Dimorphic Control of Parenting Behavior by the Medial Amygdala. Cell 176, 1206–1221 e1218. 10.1016/j.cell.2019.01.024.

9. Autry, A.E., Wu, Z., Kapoor, V., Kohl, J., Bambah-Mukku, D., Rubinstein, N.D., Marin-Rodriguez, B., Carta, I., Sedwick, V., Tang, M., and Dulac, C. (2021). Urocortin-3 neurons in the mouse perifornical area promote infant-directed neglect and aggression. Elife 10, e64680. 10.7554/eLife.64680.

10. Sato, K., Hamasaki, Y., Fukui, K., Ito, K., Miyamichi, K., Minami, M., and Amano, T. (2020). Amygdalohippocampal Area Neurons That Project to the Preoptic Area Mediate Infant-Directed Attack in Male Mice. J Neurosci 40, 3981–3994. 10.1523/JNEUROSCI.0438-19.2020.

11. Kohl, J., Babayan, B.M., Rubinstein, N.D., Autry, A.E., Marin-Rodriguez, B., Kapoor, V., Miyamishi, K., Zweifel, L.S., Luo, L., Uchida, N., and Dulac, C. (2018). Functional circuit architecture underlying parental behaviour. Nature 556, 326–331. 10.1038/s41586-018-0027-0.

12. Wu, Z., Autry, A.E., Bergan, J.F., Watabe-Uchida, M., and Dulac, C.G. (2014). Galanin neurons in the medial preoptic area govern parental behaviour. Nature 509, 325–330. 10.1038/nature13307.

13. Moffitt, J.R., Bambah-Mukku, D., Eichhorn, S.W., Vaughn, E., Shekhar, K., Perez, J.D., Rubinstein, N.D., Hao, J., Regev, A., Dulac, C., and Zhuang, X. (2018). Molecular, spatial, and functional single-cell profiling of the hypothalamic preoptic region. Science 362, eaau5324 10.1126/science.aau5324.

14. Yoshihara, C., Tokita, K., Maruyama, T., Kaneko, M., Tsuneoka, Y., Fukumitsu, K., Miyazawa, E., Shinozuka, K., Huang, A.J., Nishimori, K., et al. (2021). Calcitonin receptor signaling in the medial preoptic area enables risk-taking maternal care. Cell Rep 35, 109204. 10.1016/j.celrep.2021.109204.

15. Stagkourakis, S., Smiley, K.O., Williams, P., Kakadellis, S., Ziegler, K., Bakker, J., Brown, R.S.E., Harkany, T., Grattan, D.R., and Broberger, C. (2020). A Neuro-hormonal Circuit for Paternal Behavior Controlled by a Hypothalamic Network Oscillation. Cell 182, 960–975 e915. 10.1016/j.cell.2020.07.007.

16. Mei, L., Yan, R., Yin, L., Sullivan, R.M., and Lin, D. (2023). Antagonistic circuits mediating infanticide and maternal care in female mice. Nature 618, 1006–1016. 10.1038/s41586-023-06147-9.

17. Froemke, R.C., and Young, L.J. (2021). Oxytocin, Neural Plasticity, and Social Behavior. Annu Rev Neurosci 44, 359–381. 10.1146/annurev-neuro-102320-102847.

18. He, Z., Zhang, L., Hou, W., Zhang, X., Young, L.J., Li, L., Liu, L., Ma, H., Xun, Y., Lv, Z., et al. (2021). Paraventricular Nucleus Oxytocin Subsystems Promote Active Paternal Behaviors in Mandarin Voles. The Journal of neuroscience : the official journal of the Society for Neuroscience 41, 6699–6713. 10.1523/JNEUROSCI.2864-20.2021.

19. Inada, K., Hagihara, M., Tsujimoto, K., Abe, T., Konno, A., Hirai, H., Kiyonari, H., and Miyamichi, K. (2022). Plasticity of neural connections underlying oxytocin-mediated parental behaviors of male mice. Neuron 110, 2009–2023 e2005. 10.1016/j.neuron.2022.03.033.

20. Yoshihara, C., Numan, M., and Kuroda, K.O. (2018). Oxytocin and Parental Behaviors. Curr Top Behav Neurosci 35, 119–153. 10.1007/7854_2017_11.

21. Theofanopoulou, C., Gedman, G., Cahill, J.A., Boeckx, C., and Jarvis, E.D. (2021). Universal nomenclature for oxytocin-vasotocin ligand and receptor families. Nature 592, 747–755. 10.1038/s41586-020-03040-7.

22. Rigney, N., de Vries, G.J., Petrulis, A., and Young, L.J. (2022). Oxytocin, Vasopressin, and Social Behavior: From Neural Circuits to Clinical Opportunities. Endocrinology 163, 1–13. 10.1210/endocr/bqac111.

23. Pedersen, C.A., Ascher, J.A., Monroe, Y.L., and Prange, A.J., Jr. (1982). Oxytocin induces maternal behavior in virgin female rats. Science 216, 648–650. 10.1126/science.7071605.

24. Koshimizu, T.A., Nakamura, K., Egashira, N., Hiroyama, M., Nonoguchi, H., and Tanoue, A. (2012). Vasopressin V1a and V1b receptors: from molecules to physiological systems. Physiol Rev 92, 1813–1864. 10.1152/physrev.00035.2011.

25. Song, Z., and Albers, H.E. (2018). Cross-talk among oxytocin and arginine-vasopressin receptors: Relevance for basic and clinical studies of the brain and periphery. Front Neuroendocrinol 51, 14–24. 10.1016/j.yfrne.2017.10.004.

26. Dumais, K.M., and Veenema, A.H. (2016). Vasopressin and oxytocin receptor systems in the brain: Sex differences and sex-specific regulation of social behavior. Front Neuroendocrinol 40, 1–23. 10.1016/j.yfrne.2015.04.003.

27. Szot, P., Bale, T.L., and Dorsa, D.M. (1994). Distribution of messenger RNA for the vasopressin V1a receptor in the CNS of male and female rats. Brain Res Mol Brain Res 24, 1–10. 10.1016/0169-328x(94)90111-2.

28. Smith, C.J.W., Poehlmann, M.L., Li, S., Ratnaseelan, A.M., Bredewold, R., and Veenema, A.H. (2017). Age and sex differences in oxytocin and vasopressin V1a receptor binding densities in the rat brain: focus on the social decision-making network. Brain Struct Funct 222, 981–1006. 10.1007/s00429-016-1260-7.

29. Newmaster, K.T., Nolan, Z.T., Chon, U., Vanselow, D.J., Weit, A.R., Tabbaa, M., Hidema, S., Nishimori, K., Hammock, E.A.D., and Kim, Y. (2020). Quantitative cellular-resolution map of the oxytocin receptor in postnatally developing mouse brains. Nat Commun 11, 1885. 10.1038/s41467-020-15659-1.

30. Inada, K., Tsujimoto, K., Yoshida, M., Nishimori, K., and Miyamichi, K. (2022). Oxytocin signaling in the posterior hypothalamus prevents hyperphagic obesity in mice. Elife 11, e75718. 10.7554/eLife.75718.

31. Dolen, G., Darvishzadeh, A., Huang, K.W., and Malenka, R.C. (2013). Social reward requires coordinated activity of nucleus accumbens oxytocin and serotonin. Nature 501, 179–184. 10.1038/nature12518.

32. Hung, L.W., Neuner, S., Polepalli, J.S., Beier, K.T., Wright, M., Walsh, J.J., Lewis, E.M., Luo, L., Deisseroth, K., Dolen, G., and Malenka, R.C. (2017). Gating of social reward by oxytocin in the ventral tegmental area. Science 357, 1406–1411. 10.1126/science.aan4994.

33. Osakada, T., Yan, R., Jiang, Y., Wei, D., Tabuchi, R., Dai, B., Wang, X., Zhao, G., Wang, C.X., Liu, J.J., et al. (2024). A dedicated hypothalamic oxytocin circuit controls aversive social learning. Nature 626, 347–356. 10.1038/s41586-023-06958-w.

34. Raam, T., McAvoy, K.M., Besnard, A., Veenema, A.H., and Sahay, A. (2017). Hippocampal oxytocin receptors are necessary for discrimination of social stimuli. Nat Commun 8, 2001. 10.1038/s41467-017-02173-0.

35. Yao, S., Bergan, J., Lanjuin, A., and Dulac, C. (2017). Oxytocin signaling in the medial amygdala is required for sex discrimination of social cues. Elife 6, e31373. 10.7554/eLife.31373.

36. Wahis, J., Baudon, A., Althammer, F., Kerspern, D., Goyon, S., Hagiwara, D., Lefevre, A., Barteczko, L., Boury-Jamot, B., Bellanger, B., et al. (2021). Astrocytes mediate the effect of oxytocin in the central amygdala on neuronal activity and affective states in rodents. Nat Neurosci 24, 529–541. 10.1038/s41593-021-00800-0.

37. Marlin, B.J., Mitre, M., D’Amour J, A., Chao, M.V., and Froemke, R.C. (2015). Oxytocin enables maternal behaviour by balancing cortical inhibition. Nature 520, 499–504. 10.1038/nature14402.

38. Schiavo, J.K., Valtcheva, S., Bair-Marshall, C.J., Song, S.C., Martin, K.A., and Froemke, R.C. (2020). Innate and plastic mechanisms for maternal behaviour in auditory cortex. Nature 587, 426–431. 10.1038/s41586-020-2807-6.

39. Manning, M., Misicka, A., Olma, A., Bankowski, K., Stoev, S., Chini, B., Durroux, T., Mouillac, B., Corbani, M., and Guillon, G. (2012). Oxytocin and vasopressin agonists and antagonists as research tools and potential therapeutics. J Neuroendocrinol 24, 609–628. 10.1111/j.1365-2826.2012.02303.x.

40. Yang, C.F., Chiang, M.C., Gray, D.C., Prabhakaran, M., Alvarado, M., Juntti, S.A., Unger, E.K., Wells, J.A., and Shah, N.M. (2013). Sexually dimorphic neurons in the ventromedial hypothalamus govern mating in both sexes and aggression in males. Cell 153, 896–909. 10.1016/j.cell.2013.04.017.

41. Harris, J.A., Hirokawa, K.E., Sorensen, S.A., Gu, H., Mills, M., Ng, L.L., Bohn, P., Mortrud, M., Ouellette, B., Kidney, J., et al. (2014). Anatomical characterization of Cre driver mice for neural circuit mapping and manipulation. Front Neural Circuits 8, 76. 10.3389/fncir.2014.00076.

42. Xu, S., Yang, H., Menon, V., Lemire, A.L., Wang, L., Henry, F.E., Turaga, S.C., and Sternson, S.M. (2020). Behavioral state coding by molecularly defined paraventricular hypothalamic cell type ensembles. Science 370, eabb2494. 10.1126/science.abb2494.

43. Rigney, N., Whylings, J., de Vries, G.J., and Petrulis, A. (2021). Sex Differences in the Control of Social Investigation and Anxiety by Vasopressin Cells of the Paraventricular Nucleus of the Hypothalamus. Neuroendocrinology 111, 521–535. 10.1159/000509421.

44. Zhan, S., Qi, Z., Cai, F., Gao, Z., Xie, J., and Hu, J. (2024). Oxytocin neurons mediate stress-induced social memory impairment. Curr Biol 34, 36–45 e34. 10.1016/j.cub.2023.11.037.

45. Takayanagi, Y., Yoshida, M., Bielsky, I.F., Ross, H.E., Kawamata, M., Onaka, T., Yanagisawa, T., Kimura, T., Matzuk, M.M., Young, L.J., and Nishimori, K. (2005). Pervasive social deficits, but normal parturition, in oxytocin receptor-deficient mice. Proc Natl Acad Sci U S A 102, 16096–16101. 10.1073/pnas.0505312102.

46. Sharma, K., LeBlanc, R., Haque, M., Nishimori, K., Reid, M.M., and Teruyama, R. (2019). Sexually dimorphic oxytocin receptor-expressing neurons in the preoptic area of the mouse brain. PLoS One 14, e0219784. 10.1371/journal.pone.0219784.

47. Mitre, M., Marlin, B.J., Schiavo, J.K., Morina, E., Norden, S.E., Hackett, T.A., Aoki, C.J., Chao, M.V., and Froemke, R.C. (2016). A Distributed Network for Social Cognition Enriched for Oxytocin Receptors. J Neurosci 36, 2517–2535. 10.1523/JNEUROSCI.2409-15.2016.

48. Rood, B.D., and De Vries, G.J. (2011). Vasopressin innervation of the mouse (Mus musculus) brain and spinal cord. J Comp Neurol 519, 2434–2474. 10.1002/cne.22635.

49. Miyamichi, K., Shlomai-Fuchs, Y., Shu, M., Weissbourd, B.C., Luo, L., and Mizrahi, A. (2013). Dissecting local circuits: parvalbumin interneurons underlie broad feedback control of olfactory bulb output. Neuron 80, 1232–1245. 10.1016/j.neuron.2013.08.027.

50. Inoue, Y.U., Miwa, H., Hori, K., Kaneko, R., Morimoto, Y., Koike, E., Asami, J., Kamijo, S., Yamada, M., Hoshino, M., and Inoue, T. (2022). Targeting Neurons with Functional Oxytocin Receptors: A Novel Set of Simple Knock-In Mouse Lines for Oxytocin Receptor Visualization and Manipulation. eNeuro 9. 10.1523/ENEURO.0423-21.2022.

51. Bosch, O.J., Pfortsch, J., Beiderbeck, D.I., Landgraf, R., and Neumann, I.D. (2010). Maternal behaviour is associated with vasopressin release in the medial preoptic area and bed nucleus of the stria terminalis in the rat. J Neuroendocrinol 22, 420–429. 10.1111/j.1365-2826.2010.01984.x.

52. Waltenspuhl, Y., Ehrenmann, J., Vacca, S., Thom, C., Medalia, O., and Pluckthun, A. (2022). Structural basis for the activation and ligand recognition of the human oxytocin receptor. Nat Commun 13, 4153. 10.1038/s41467-022-31325-0.

53. Kawamata, M., Mitsui-Saito, M., Kimura, T., Takayanagi, Y., Yanagisawa, T., and Nishimori, K. (2003). Vasopressin-induced contraction of uterus is mediated solely by the oxytocin receptor in mice, but not in humans. Eur J Pharmacol 472, 229–234. 10.1016/s0014-2999(03)01914-9.

54. Schorscher-Petcu, A., Sotocinal, S., Ciura, S., Dupre, A., Ritchie, J., Sorge, R.E., Crawley, J.N., Hu, S.B., Nishimori, K., Young, L.J., et al. (2010). Oxytocin-induced analgesia and scratching are mediated by the vasopressin-1A receptor in the mouse. J Neurosci 30, 8274–8284. 10.1523/JNEUROSCI.1594-10.2010.

55. Qian, T., Wang, H., Wang, P., Geng, L., Mei, L., Osakada, T., Wang, L., Tang, Y., Kania, A., Grinevich, V., et al. (2023). A genetically encoded sensor measures temporal oxytocin release from different neuronal compartments. Nat Biotechnol 41, 944–957. 10.1038/s41587-022-01561-2.

56. Ammari, R., Monaca, F., Cao, M., Nassar, E., Wai, P., Del Grosso, N.A., Lee, M., Borak, N., Schneider-Luftman, D., and Kohl, J. (2023). Hormone-mediated neural remodeling orchestrates parenting onset during pregnancy. Science 382, 76–81. 10.1126/science.adi0576.

57. Yaguchi, K., Hagihara, M., Konno, A., Hirai, H., Yukinaga, H., and Miyamichi, K. (2023). Dynamic modulation of pulsatile activities of oxytocin neurons in lactating wild-type mice. PLoS One 18, e0285589. 10.1371/journal.pone.0285589.

58. Lein, E.S., Hawrylycz, M.J., Ao, N., Ayres, M., Bensinger, A., Bernard, A., Boe, A.F., Boguski, M.S., Brockway, K.S., Byrnes, E.J., et al. (2007). Genome-wide atlas of gene expression in the adult mouse brain. Nature 445, 168–176. 10.1038/nature05453.

59. Son, S.J., Filosa, J.A., Potapenko, E.S., Biancardi, V.C., Zheng, H., Patel, K.P., Tobin, V.A., Ludwig, M., and Stern, J.E. (2013). Dendritic peptide release mediates interpopulation crosstalk between neurosecretory and preautonomic networks. Neuron 78, 1036–1049. 10.1016/j.neuron.2013.04.025.

60. Osakada, F., and Callaway, E.M. (2013). Design and generation of recombinant rabies virus vectors. Nature protocols 8, 1583–1601. 10.1038/nprot.2013.094.

61. Ishii, K.K., Osakada, T., Mori, H., Miyasaka, N., Yoshihara, Y., Miyamichi, K., and Touhara, K. (2017). A Labeled-Line Neural Circuit for Pheromone-Mediated Sexual Behaviors in Mice. Neuron 95, 123–137 e128. 10.1016/j.neuron.2017.05.038.

